# Transient evolutionary epidemiology of viral adaptation and lethal mutagenesis

**DOI:** 10.1101/2024.05.14.594065

**Authors:** Martin Guillemet, Erwan Hardy, Denis Roze, Sylvain Gandon

## Abstract

Beneficial mutations drive the within-host adaptation of viral populations and can prolong the duration of host infection. Yet, most mutations are not adaptive and the increase of the mean fitness of viral populations is hampered by deleterious and lethal mutations. Because of this ambivalent role of mutations, it is unclear if a higher mutation rate boosts or slows down viral adaptation. Here we study the interplay between selection, mutation, genetic drift and within-host dynamics of viral populations. We obtain good approximations for the transient evolutionary epidemiology of viral adaptation under the assumption that the mutation rate is high and the effects of non-lethal mutations remain small. We use this theoretical framework to discuss the feasibility of lethal mutagenesis to treat viral infections in the light of quantitative predictions we obtained for the critical mutation rates of a range of different viruses.

## Introduction

The within-host dynamics of viral infections depends both on the availability of susceptible host cells and the ability of the virus to infect and exploit these cells. Mathematical epidemiology provides a theoretical framework to model how these life-history traits affect the dynamics of viral populations [1–3]. Yet these traits are not constant but may evolve and change during the course of the host infection. Indeed, many viruses undergo high mutation rates [4], yielding substantial genetic and phenotypic diversity of within-host viral populations. This influx of mutations challenges the simplicity of classical models of viral dynamics and has led to the concept of *quasispecies* to describe the dynamics of viruses with high mutation rates [5, 6]. The effects of high mutation rates can also be captured within the classical population genetics framework [7–9]. As most mutations have deleterious effects, the constant influx of mutations generates a *mutation load* which measures the difference between fitness of the fittest strain and the mean fitness of the population [10]. In fact, some mutations can prevent viral replication and can be considered as *lethal mutations* [11]. The massive impact of deleterious mutations on viral fitness led to the “lethal mutagenesis hypothesis” which states that there is a mutation rate above which a viral population cannot grow and is driven to extinction [8, 12]. Drugs increasing mutation rates may thus constitute a broadly applicable therapeutic strategy against many viruses [13, 14], including SARS-CoV-2 [15–19]. A better evaluation of the balance between the therapeutic potential and the risk associated with these drugs relies on a better understanding of the within-host viral dynamics with increased mutation rates.

First, it is important to realise that mutation is a double-edged sword: even if most mutations are deleterious, some may be beneficial and could speed up within-host evolution. Some concern has emerged regarding the potential risk of “sublethal mutagenesis” that may result in immune escape or higher transmission [20–23]. Experimental evolution of bacteriophage T7 under high mutation rates showed that the exposure to a mutagen may boost adaptation and increase the mean fitness of the viral population [12, 24]. Several theoretical models explored the balance between the effects of deleterious and beneficial mutations [9, 25–27]. Yet, another aspect which is often overlooked in previous models is the epidemiological dynamics taking place within the infected host. The drop of the mean fitness of the virus consecutive to accumulation of deleterious mutations is expected to reduce the within-host viral load. This may trigger a rebound in the density of susceptible host cells, higher viral replication and, eventually, treatment failure. Hence, the critical mutation rate above which the virus population is expected to go extinct needs to account for this demographic feedback [9].

Second, even if lethal mutagenesis is a deterministic process occurring when the mutation load becomes overwhelmingly high, the effect of higher mutation rates may be amplified by demographic stochasticity. In finite populations, the most fit, least-loaded genotype will be lost as genetic drift overwhelms the effect of natural selection. This will result in a lower population size, thus increasing the magnitude of genetic drift and the drop of mean fitness. There is thus a synergy between the demographic and evolutionary dynamics [28–30]. However, even a small influx of compensatory mutations can halt the meltdown of small populations and prevent their extinction [31]. But demographic stochasticity could increase the risk of extinction when the viral population becomes transiently very low after the start of a mutagenic treatment [27]. Yet, the theory of lethal mutagenesis is based on the analysis of the equilibrium of the deterministic dynamics of viral populations and the identification of the critical mutation rate beyond which the virus is driven to extinction. Yet, we currently lack a good understanding of the effects of higher mutation rates on the transient within-host dynamics of viral adaptation.

In the present work, we study the joint epidemiological and evolutionary within-host dynamics of a viral population exposed to high mutation rates. We use Fisher’s Geometric Model (FGM) to link the phenotype of the virus to its within-host transmission rate [9]. This geometric model of adaptation generates distributions of fitness effects of mutations and allows us to account for deleterious, (nearly) neutral or beneficial effects of mutations [32, 33]. Because these fitness distributions depend on the parental genotype, the model accounts also for pervasive epistasis in fitness between mutations [34]. In addition, we account for a distinct type of strictly lethal mutations. The analysis of the long-term equilibrium of the epidemiology and evolution of viral dynamics allows us to identify the critical mutation rate leading to the within-host eradication of the virus [9]. We estimate these critical mutation rates for different viruses using previously measured parameters, which provide an estimate of the required efficacy of mutagenic drug to lead to lethal mutagenesis. Crucially, we derive approximations for the transient within-host evolutionary epidemiology of the virus. The analysis of these transient dynamics allows us to identify the pros and cons of higher mutation rates under different scenarios. In particular, we discuss the potential risks of lethal mutagenesis for the treatment of viral infections. Finally, we check the validity of our deterministic predictions with stochastic simulations.

## 1 Model

We assume that different strains of the virus may circulate within the host. To describe this diversity, we follow the distribution of the different phenotypes associated with these strains. In this study we consider that transmission is the only *life-history trait* under selection and we assume that it is governed by *n* underlying *phenotypic traits*. We define **x** as a vector of size *n*, where each dimension refers to an independent continuous trait. To model the dependence of the transmission rate *β*_**x**_ on **x**, we use a fitness landscape based on a quadratic Fisher’s geometric model where the transmission rate depends on the Euclidian distance of the vector **x** to the optimum (at the origin i.e. at **x** = (0, 0, .., 0)) such that :

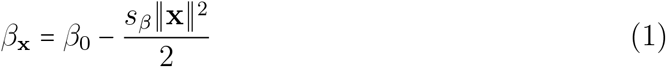

where ∥**x**∥ ^2^ is the squared norm of the phenotypic vector **x** (**Figure 1**). The term *s*_*β*_ governs the curvature of the fitness landscape and represents the strength of the directional selection towards the optimum where the transmission rate is *β*_0_. Note that we use this quadratic form to approximate a Gaussian shape of the fitness landscape [9] and the division by 2 is for consistency with earlier work [35]. Indeed, equation (1) can yield a good approximation when the virus is not too far from the optimum. However, note that this approximation breaks down when the phenotype is very far from the optimum as transmission rate can become negative. We show how this fitness landscape links the *n* underlying phenotypes with the pathogens’ life-history trait (i.e. transmission rate *β*) in **Figure 1**.

**Figure 1:**
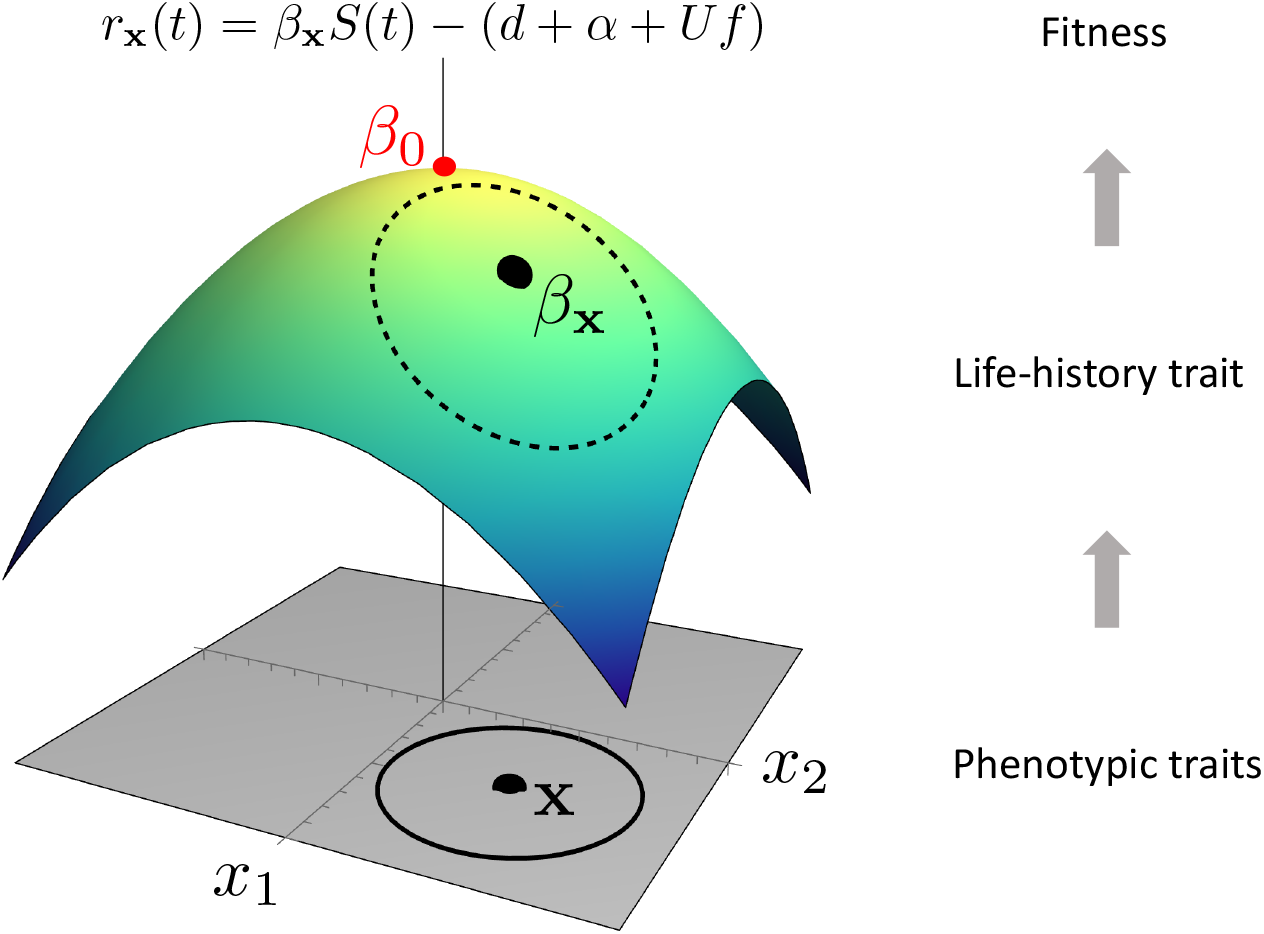
The fitness landscape links the underlying phenotypic traits with life-history traits and malthusian fitness. We represent our phenotype to life-history trait landscape in two dimensions (*n=* 2). A phenotype **x** is translated to a transmission rate *β*_**x**_ using equation (1). This life-history trait is translated to a fitness value with equation (5), which depends on time through the number of susceptible cells *S* (*t*). A black circle is shown around phenotype **x**, which is translated to a dashed red circle of transmission rates, showing how the FGM distorts the distributions.

In the following, we will track the evolution of the distribution of the life-history trait *β* itself. In particular, the mean transmission rate can be written in the following way (see equation (S.2)):

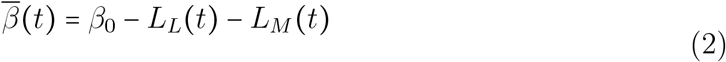

as a function of the “lag load”, *L*_*L*_(*t*), and the “mutation load”, *L*_*M*_(*t*)[36]:

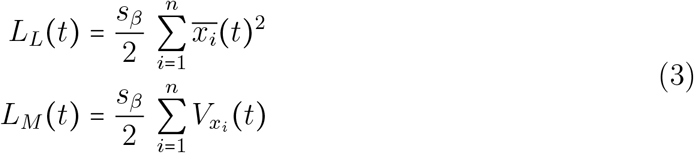

The lag load depends on the sum of the distance to the optimum 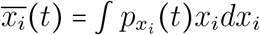 on each dimension *i* ∈ [1, *n*], where 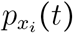 refers to the proportion of the virus population with phenotypic trait *x*_*i*_. The mutation load depends on the sum of the phenotypic variance 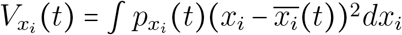 on each dimension *i* ∈ [1, *n*]. These load terms depend on the number of dimensions *n* because of the so called “cost of complexity” [32, 33]. A lower strength of selection *s*_*β*_ (a measure of the steepness of the landscape around the optimum) decreases both the lag load and the mutation load, as it makes the fitness landscape flatter.

To model the joint epidemiological and evolutionary within-host dynamics of a virus population we assume that the viral pathogen has access to a density *S* of susceptible host cells. We focus on the dynamics of the density *I* of infected cells and we do not explicitly model the dynamics of the free virus stage because we assume the lifetime of an infected cell is much longer than a free virus particle [3, 9]. Host cells are produced at a constant rate *b* and die at a constant rate *d. I*_**x**_ (*t*) refers to the density of host cells infected by strain **x**. An infected cell of phenotype **x** infects susceptible cells with a transmission rate *β*_**x**_ and dies at a rate (*d*+ *α* +*Uf)* where *α* is the virulence and *Uf* is the rate of lethal mutations. One can use the ratio *U* (1− *f*)/(*d*+ *α* +*Uf*) to get the expected number of non-lethal mutations arising in an infected cell from the the time it is infected to its death. These lethal mutations produce cells that are infected with a non-transmissible form of the virus. As there is no density-dependent reproduction of susceptible host cells in our model, the cells infected by viruses with lethal mutations do not affect the dynamics of the system. Consequently, we do not monitor the density of the cells infected by non-transmissible pathogens, and we simply treat lethal mutations as an additional death term. We assume that each host cell can only be infected with a single viral strain (i.e. no multiple infections). The overall death rate of the infected cells is not affected by the phenotype of the virus and the above life cycle yields the following system of ordinary differential equations (the upper dot represents time derivation):

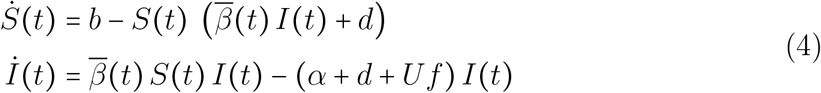

where *I (t)* =∫*I*_x_ (*t*) *d*^*n*^**x** is the total density of infected cells (i.e. cells infected by a transmissible virus), 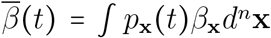 is the mean transmission rate and *p*_x_ (*t*)= *I*_**x**_ (*t*)/ *I*(*t* is the frequency of the phenotype **x** in the infected population. We can now introduce the per capita growth rate (i.e. Malthusian fitness) of the phenotype **x** and the mean growth rate of the pathogen (i.e. the mean fitness) :

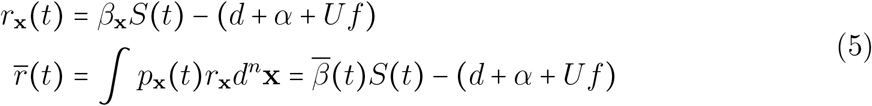

The sign of the mean fitness indicates whether a population of infected cells can invade a population of hosts of size *S*(*t*) (i.e.,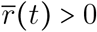) or if it drops and eventually goes extinct (i.e.,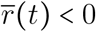).

Next we need to describe the dynamics of the density *I*_**x**_ (*t*) of cells infected by each phenotype **x** by introducing *U* the rate of mutation of the virus. This mutation process is assumed to be constant and unconditional on a transmission event. With probability *f* the mutation is lethal and we model this effect as an additional rate of mortality *Uf* for the infected cells. With probability 1− *f* the mutation is non-lethal and the new phenotype becomes **x+ u**, where the mutation effect **u** is sampled in an isotropic multivariate normal distribution *ρ (***u**) with mean 0 and variance *λ*. This mutation process yields the following integro-differential equation:

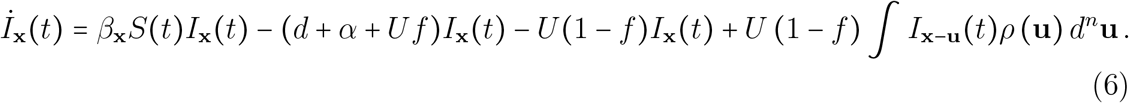

The mutational variance *λ* is easily interpreted as it directly relates to the the mean effect of random mutations on transmission rate *µ*_*β*_ :

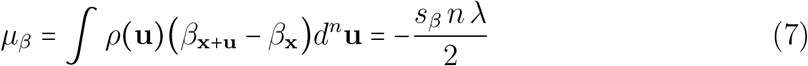

We can directly relate this to the mean effect of mutations on fitness *µ*_*r*_ (*t*) = *µ*_*β*_*S* (*t*) which depends on the density of susceptible cells at time *t*, but does not depend on the parental phenotype **x**. Note that, since *µ*_*β<*_ 0, the mean effect of mutations on fitness is always negative, meaning that mutated strains are always, on average, worse than their parental strain.

In the following, we use different approaches to monitor the epidemiological and evolutionary dynamics described in (6). First, for the sake of simplicity, we use a Weak Selection Strong Mutation (WSSM) approximation which implies that adaptation is the result of many mutations of small effects. In this regime, the distribution of phenotypes remains Gaussian [37]. Second, we relax this assumption and we study how mutations of larger effects can influence the evolutionary dynamics of the virus using a moment closure approximation. Third, we check the robustness of our approximations using numerical simulations, where we study the dynamics of discrete phenotypes on a 2D grid, with or without the assumption of small mutational effects. Finally, we use our theoretical framework to discuss the feasibility and the risks associated with lethal mutagenesis therapy under different scenarios. In particular, we use stochastic simulations to explore the effect of finite host and viral population sizes on our theoretical predictions on the probability of treatment failure.

## 2 Results

We use equation (6) to obtain dynamical equations for the cumulants of the distribution of the phenotypes **x**. Under the WSSM assumptions, we can neglect higher order terms of the mutational variance (i.e., *λ*^2^, *λ*^3^ etc.), the phenotypic distribution is Gaussian and the evolutionary dynamics can be described with the first two moments of the phenotypic distribution [35]. Because of the isometry of the fitness landscape and of the model of mutation, the phenotypic distribution is the same along all dimensions and we can fully capture the evolutionary dynamics on a single dimension *x*, which yields (Supplementary Information S2.5):

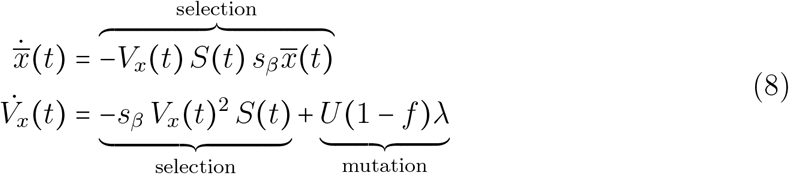

Note how the epidemiological dynamics (4) feeds back on the evolution of the phenotypic distribution. The first equation shows how the mean phenotype moves towards the optimum at a speed governed by (i) the amount of susceptible cells *S* (*t*), (ii) the phenotype variance *V*_*x*_ (*t*) and (iii) the mean distance to the optimum 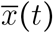. The dynamics of *V*_*x*_ (*t*) results from the balance between the effect of natural selection which consumes this variation, and the effect of mutation which introduces more genetic variation. Interestingly, the number of dimensions *n* only appears in the epidemiological equations through the load terms defined which yields (see S.35):

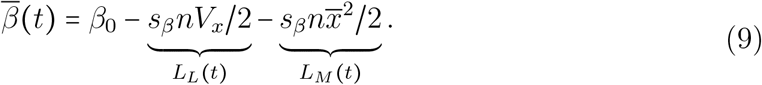

Using the assumption that the phenotypic distribution remains Gaussian we obtain a dynamical equation for the mean transmission rate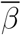:

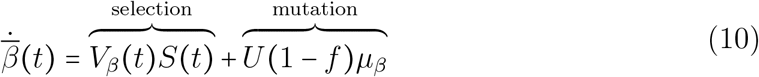

where *V*_*β*_ (*t*) *= V*_*x*_ (*t*) *s*_*β*_ (2*L*_*L*_ (*t*) + *L*_*M*_ (*t*)) is the variance in transmission rate. The mean transmission rate is increased by natural selection with a speed controlled by the variance in transmission rate, scaled by the density of susceptible cells. In contrast, the direct effect of the mutation process is negative and equals to the rate of non-lethal mutations times the mean effect of these mutations on transmission rate. Indeed, as discussed above, the expected effect on the life history-trait is deleterious (see equation (7)).

### 2.1 Evolutionary equilibrium and critical mutation rate

Next, we focus on the long-term dynamics of the viral population to determine what is the critical mutation rate *U*_*c*_ above which natural selection is overwhelmed by the accumulation of deleterious mutations and the viral population is driven to extinction. At equilibrium, the mean phenotype is at the optimum and 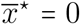 (where the indicates that the system is at equilibrium). The phenotypic distribution of this long-term equilibrium depends both on the epidemiology (equation (4)) and the evolution of the virus (equations (8) and (10)) yielding :

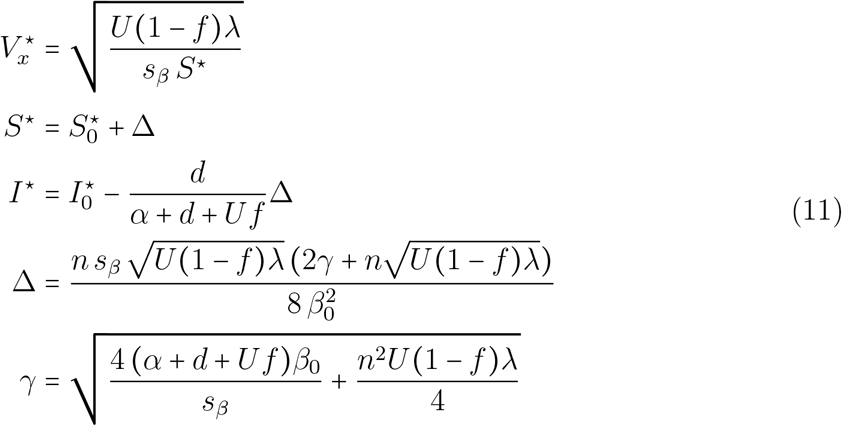

where 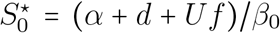 and 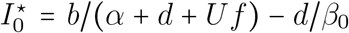 are the equilibrium densities of the susceptible and infected cells, respectively, when the virus population is monomorphic and the virus is at the optimum (i.e. *β= β*_0_ is maximal). As expected, equation (11) shows that the constant influx of non-lethal mutations introduces a mutation load 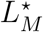 that leads to a reduced density of infected cells at equilibrium (and an increased density of susceptible cells) because Δ> 0. Using (9) we can compute the mean transmission rate 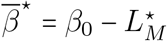.

The critical mutation rate *U*_*c*_ is reached when 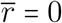 and can be obtained from (11) (see Supplementary Information S2.8) by finding the value of the mutation rate where *I*^⋆^ = 0 which yields (see also [9]):

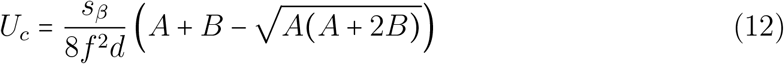

with *A=(* 1 −*f) λb n*^2^ and *B* = 8*f (β*_0_ *b*− *d(d* +*α))/ s*_*β*_. As expected, the fraction *f* of lethal mutations has a massive influence on the critical mutation rate:

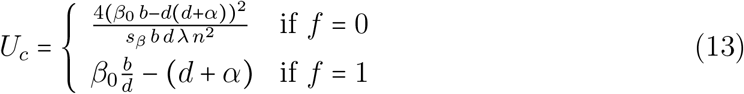

To understand the expression of the critical mutation rate it is useful to note that *U*_*c*_ is also the mutation rate above which the virus population cannot grow when the density of susceptible cells is *b/ d* (i.e., the equilibrium density of the host cells in the absence of the virus). Using equation (5), (2), (9) and (11) to obtain the mean fitness of the population at equilibrium when *S*(*t*) = *b*/*d*, yields the following condition:

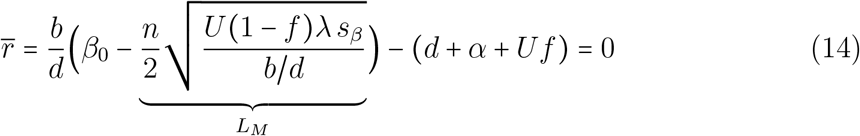

The final term in (14) accounts for the direct effect of lethal mutations: increasing the proportion of lethal mutations increases the *death rate* of infections, and consequently decreases the critical mutation rate leading to viral extinctions. Yet, lethal mutations have an additional effect on the *birth rate* of infections. Indeed, the first term in equation (14) refers to the rate of new infections and this rate drops with higher mutation load *L*_*M*_ (see equation (9) and (11)). This mutation load drops when most mutations are lethal because only viable mutations are accounted for in this load. Hence, increasing the proportion of lethal mutations decreases the mutation load. This effect is relatively small when the number *n* of phenotypic dimensions is low, but it can counteract the direct effect of lethal mutations when *n* becomes large. In other words, we find that the effect of the cost of complexity, which is expected to decrease the critical mutation rate for larger values of *n*, is actually dependent on the proportion of lethal mutations *f* (**Figure 2**).

**Figure 2:**
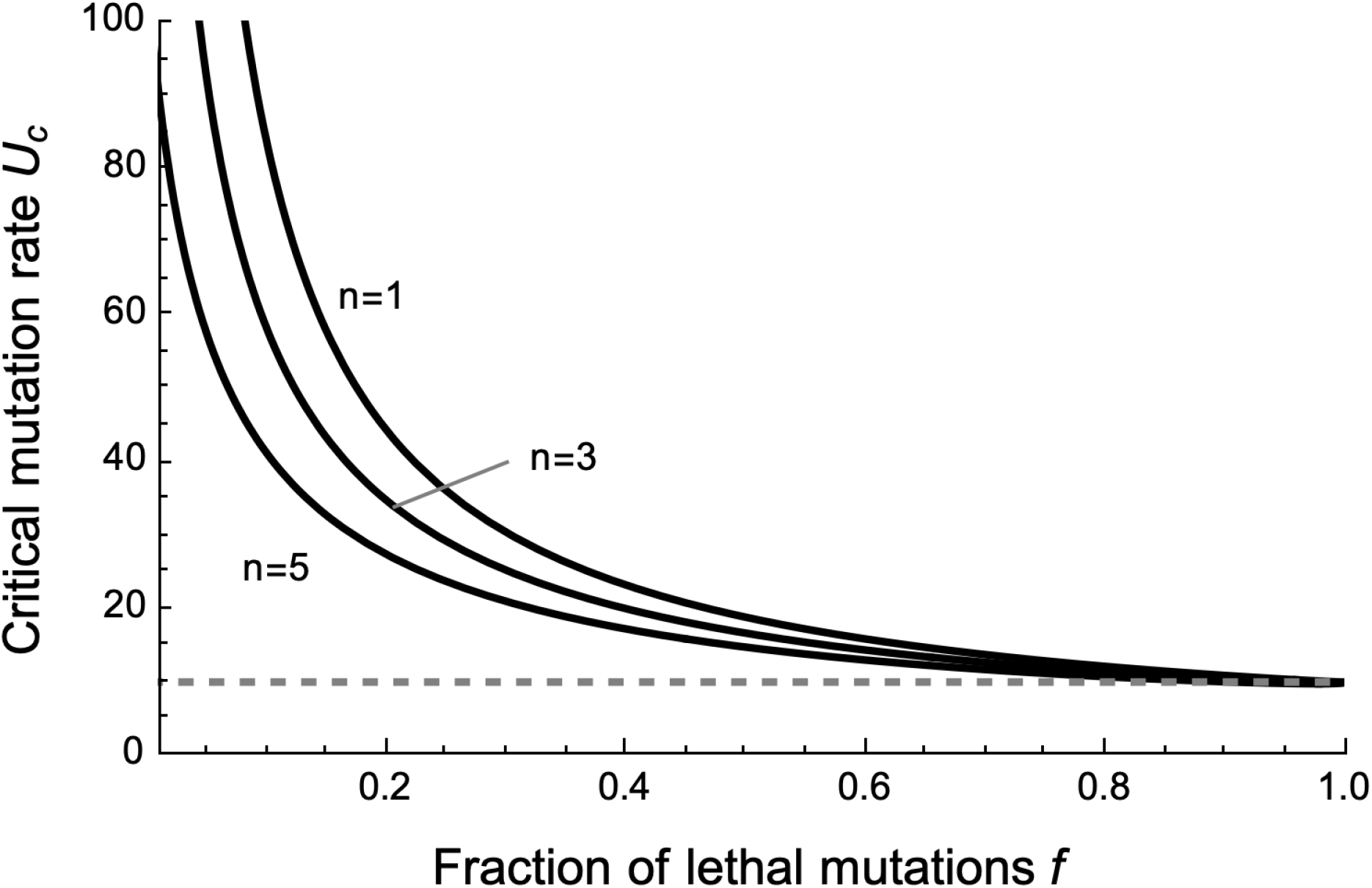
Effect of the proportion of lethal mutations and the phenotypic dimension of the fitness landscape on the critical mutation rate. The plot shows the effect of both fraction of lethal mutations *f* and number of dimensions *n* on the critical mutation rate required for lethal mutagenesis. The solid lines correspond to the critical mutation rate computed for the WSSM regime. In comparison, the dashed grey line shows the critical mutation rate predicted by the House of Cards (HC) regime, which does not vary with *f*. Parameters used: *b* = 3, *d* = 1, *β*_0_ = 4, *α* = 1, *s*_*β*_ = 1, *λ* = 0.05.

### 2.2 Critical mutation rate when selection is stronger

Under the above WSSM assumption, we neglected higher order terms of the mutational variance (i.e., *λ*^2^, see Supplementary Information). In the following we relax the weak selection assumption and these higher order terms cannot be neglected anymore. In this situation, the phenotypic distribution is no longer multivariate Gaussian. We thus have to account for the effects of *K*_3_ and *K*_4_, the third and fourth cumulants of the phenotypic distribution at equilibrium (where the ⍟ indicates this new equilibrium) and we derive in Supplementary Information S2.6:

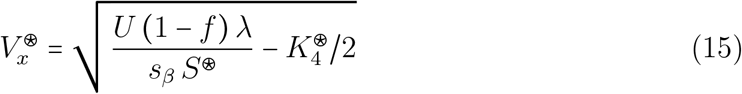

The fourth cumulant builds up with the second moment of the distribution of mutational effects *λ*^2^ and is always positive (see Supplementary Information S2.5 and Figure **Figure S1**), and the equilibrium variance is smaller than expected in the WSSM approximation (**Figure S2**). Consequently, when the effects of the mutation are stronger we need to account for *K*_4_ to compute the critical mutation rate (orange dashed line in **Figure 3**). The critical mutation rate drops with higher values of *λ* (see **Figure S3**) but the above analysis breaks down when *λ* becomes too high relative to the standing variance (see Supplementary Information).

**Figure 3:**
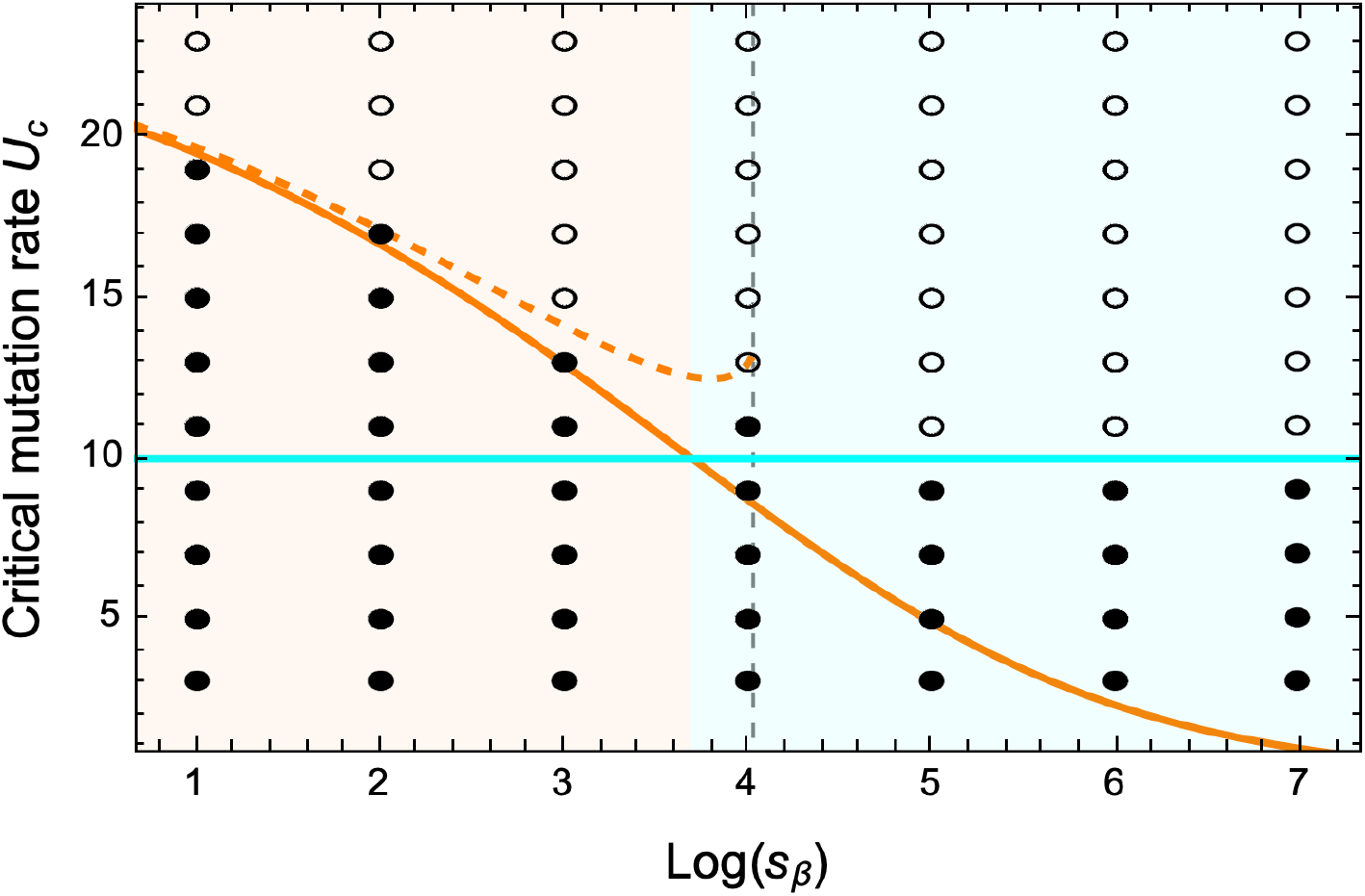
Lethal mutagenesis is feasible for lower mutation rates with steeper fitness landscapes. The critical mutation rate *U*_*c*_ is shown as a function of the selection parameter *s*_*β*_. The results are shown under three approximation: the WSSM approximation (orange), the approximation accounting for *K*_3_ and *K*_4_ cumulants (dashed orange), the House of Cards (HC) approximation (cyan). Data from numerical simulations are shown as filled circles when the infected population survives, and empty circles when the infected population goes to extinction. The shaded orange and cyan areas indicate the validity conditions of the WSSM and the House of Cards regimes, respectively (see equation (17)). The vertical dashed line shows the maximum value of *s*_*β*_ for which the WSSM+cumulant approach is valid. Parameters used: *b*= 3, *d*= 1, *β*_0_ =4, *α*= 1, *f* =0.4, *λ* =0.05, *n*= 2. Note that we use *n* = 2 here to reduce simulation time. For the stochastic simulations, we used *τ* = 0.05, *t*_*max*_ = 50

For very large values of *λ* we can use another approximation to describe the viral dynamics under a regime of mutation where the variance of mutation overwhelms the effect of the parental strain: the classical “House-of-Cards” (HC) approximation which is based on the assumption that the variance of the mutation distribution is much larger than the equilibrium variance. This approximation has been used to derive the equilibrium mutation load for haploids [35, 37]. After incorporating the influence of epidemiological feedbacks for pathogens we obtain (see Supplementary Information S2.7) the following expectation for the phenotypic variance at equilibrium:

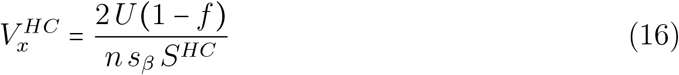

where *S*^*HC*^ is the equilibrium density of susceptible cells under the HC approximation. Note that the mutation load given by (9) becomes *L*_*M*_*= U (*1− *f*)/*S*^*HC*^ and, as discussed in [37], the mutation load on *fitness* (see equation (5) to express 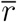 as a function of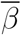) is simply equal to *U (*1− *f)*, the non-lethal mutation rate. Another striking feature of the HC approximation is the independence of equilibrium phenotypic variance to the mutational variance *λ*. Note, however, that higher order HC approximations for the equilibrium phenotypic variance do depend on *λ*^2^ [35]. Another interesting feature of the equilibrium distribution of fitness under this regime is the emergence of a spike in this distribution (i.e. a Dirac delta function) when *n>* 2, which is due to the proportion of the population having exactly the optimal fitness value [37, 38].

Numerical computations of the equilibrium phenotypic variance confirm the validity of the above approximations (**Figure S2**). As expected, the equilibrium variance predicted with the WSSM approximation is very accurate when the strength of selection is low, but this approximation breaks down when the effect of mutations becomes high. Using 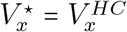,we find that the HC approximation becomes better than the WSSM approximation when:

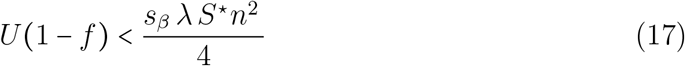

This corresponds to an upper bound on the rate of non-lethal mutations. We show in the different regimes of mutation how the critical mutation rate changes with the strength of selection in **Figure 3**. The critical mutation rate predicted by the WSSM approximation is accurate until the condition (17) is verified. For higher values of *s*_*β*_, the mutation rate required to achieve lethal mutagenesis is higher under the HC approximation than under the WSSM approximation (**Figure 3**). Interestingly, the critical mutation rate predicted under the HC approximation is the same as the one obtained under the WSSM approximation if all mutations were lethal (see (13)):

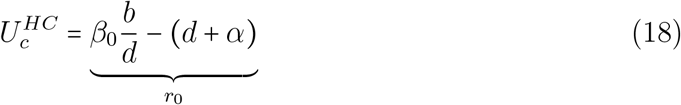

where *r*_0_ is the maximal growth rate of the optimal phenotype in a fully susceptible population. In other words, the critical mutation rate under the HC approximation depends only on the demographic parameters but not on parameters governing the fitness landscape (**Figure 3**).

We show in Supplementary Information S2.9 how the demographic parameters like *r*_0_ and the parameters characterizing the fitness landscape (*n* and the product *s*_*β*_*λS*_⋆_) and the mutation process (*U* and *f*) can be estimated experimentally. We use this approach to predict the critical mutation rates of six different viruses in **Table 1**.

**Table 1:** Critical mutation rates for six different viruses. The first 2 lines of the table correspond to observed genome sizes (kb) and mutation rates in s/n/c (substitutions per nucleotide per cell infection). These two quantities allow us to obtain the genomic mutation rate *U* in *s /c* (substitution per cell infection). The following 5 lines present parameters of demography and mutational effects: *r*_0_ is the growth rate per hour of a wildtype virus (here we assume that the wildtype is at the optimum), *E (s*_*v*_) is the mean effect of non-lethal mutations scaled by the growth rate, *V ar (s*_*v*_) is the variance of these mutational effects, *f* is the proportion of lethal mutations and *τ* is the generation time (we assume *τ* to be equal to *c*, the duration of a cell infection). We show in **Supplementary Information S2.9** how to use these measures to compute *λs*_*r*_ and *n* and the critical mutation rates corresponding to the WSSM and the HC approximations. These rates as expressed as a rate of genomic mutation per hour /*h* (or per generation *s c*). Finally, we compute the fold-increase in mutation rate required to achieve viral extinction under the WSSM and HC approximations. Gray cells refer to data from published studies [4, 49–52] and are described in the **Supplementary Information S2.9**. White cells refer to predictions derived from our model and the data in the grey cells. Values noted with a star (⋆) for SARS-CoV-2 are the average of the values obtained for the five other viruses due to the current absence of experimental data on the distribution of fitness effects.

| | Qβ | Φx174 | F1/M13 | VSV | Influenza | SARS-CoV-2 |
| --- | --- | --- | --- | --- | --- | --- |
| Genome size | 4.22 kb | 5.39 kb | 6.4 kb | 11.2 kb | 13.6 kb | 29.9 kb |
| Mutation rate | $1.4 \times 10^{-4} s/n/c$ | $1.1 \times 10^{-6} s/n/c$ | $7.8 \times 10^{-7} s/n/c$ | $3.5 \times 10^{-5} s/n/c$ | $2.3 \times 10^{-5} s/n/c$ | $1.5 \times 10^{-6} s/n/c$ |
| Genomic mutation rate $U$ | $5.91 \times 10^{-1} s/c$ | $5.93 \times 10^{-3} s/c$ | $5.00 \times 10^{-3} s/c$ | $3.92 \times 10^{-1} s/c$ | $3.13 \times 10^{-1} s/c$ | $4.49 \times 10^{-2} s/c$ |
| $r_0$ | 3.6/h | 10/h | 4.3/h | 0.9/h | 0.32/h | 0.12/h |
| $E(s_v)$ | 0.10 | 0.13 | 0.10 | 0.13 | 0.12 | 0.12* |
| $Var(s_v)$ | 0.02 | 0.05 | 0.04 | 0.04 | 0.03 | 0.04* |
| $f$ | 29% | 18% | 20% | 39% | 30% | 27%* |
| Generation time $\tau$ | 2.07h | 0.46h | 1h | 7.68h | 6-31.6h | 20.2h |
| $\lambda s_r$ | 0.61/h | 3.82/h | 1.53/h | 0.22/h | 0.070/h | 0.036/h |
| $n$ | 1.18 | 0.68 | 0.5 | 0.84 | 1.14 | 0.75 |
| $U_c^{WSSM}$ | 8.51/h<br>17.6 s/c | 35.6/h<br>16.4 s/c | 15.4/h<br>15.4 s/c | 1.50/h<br>11.5 s/c | 0.72/h<br>4.3-22.7 s/c | 0.30/h<br>6.07 s/c |
| **Fold-increase $U_c^{WSSM}/U$** | **30** | **2770** | **3080** | **29** | **14-73** | **135** |
| $U_c^{HC}$ | 3.6/h<br>7.45 s/c | 10 /h<br>4.6 s/c | 4.3/h<br>4.3 s/c | 0.8/h<br>6.1 s/c | 0.32/h<br>1.92-10.1 s/c | 0.11/h<br>2.32 s/c |
| **Fold-increase $U_c^{HC}/U$** | **13** | **776** | **860** | **16** | **6.1-32** | **51** |

### 2.3 Transient dynamics of adaptation

Next, we jointly use equations (4), (2), (8) and (10) to study the interplay between the transient within-host dynamics of the virus and the adaptation of the viral populations in the WSSM regime. In **Figure 4** we explore the dynamics of the mean transmission rate 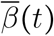 for different initial conditions. We vary the initial distance of the mean phenotype to the the optimal phenotype, and the standing variance. We contrast a scenario where the virus population is monomorphic (*V*_*x*_ (0)= 0, full line) and a scenario with some standing genetic variance (*V*_*x*_ (0) = 0.1, dashed line). As expected, regardless of these initial conditions, the dynamics converge to the same equilibrium, which is given by (2), (9) and (11). However, the initial conditions govern the speed at which this equilibrium is reached. First, the standing genetic variance induces a mutation load, which explains the lower value of 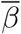 at *t* = 0 in **Figure 4**. Second, the absence of a genetic variation in the clonal population implies that the speed of adaptation is initially very slow. In fact, the mean transmission rate initially drops because of the effect of deleterious mutations (see equation (10)). Genetic variation first needs to build up before selection can act on the mean transmission rate. In contrast, the speed of adaptation is faster with standing genetic variance. This faster adaptation allows the population to rapidly overcome the initial mutation load and the mean transmission rate 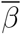 becomes rapidly higher than in initially clonal populations. Finally, the rate of change in mean transmission rate is increased when the initial distance to the optimum (or initial lag load) is higher (see equation (8)). This is due to a faster decrease in the lag load but note that it does not affect the dynamics of the phenotypic variance and thus mutation load.

**Figure 4:**
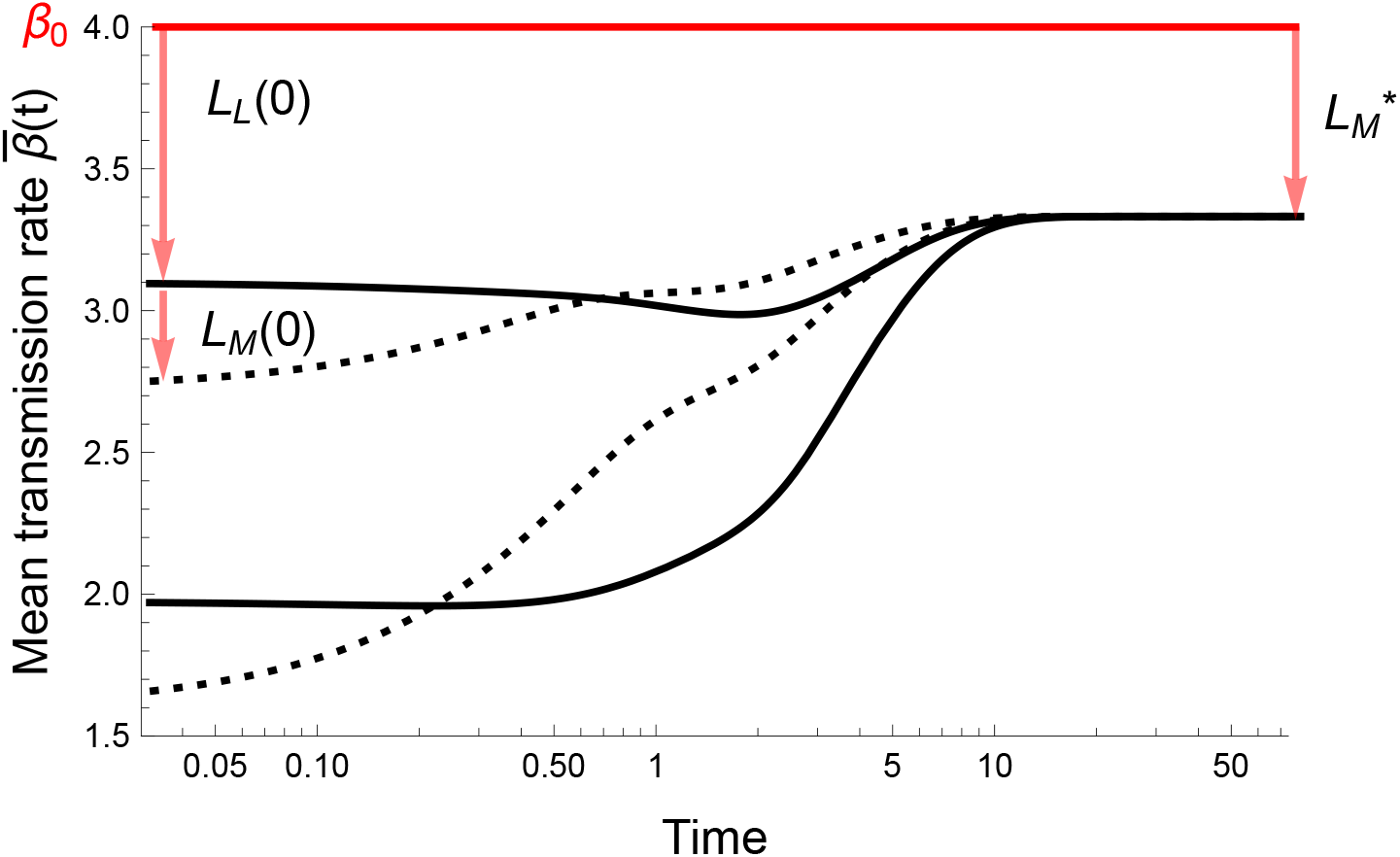
The transient evolutionary dynamics depends on the distance to the optimum and the initial phenotypic variance. We show the dynamics of mean transmission rate 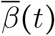 through time starting from a clonal populations at different initial values of 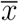.Represented in solid black the infected population is initially clonal with *V*_*x*_ (0) =0 while in dashed black the population is initially diverse with *V*_*x*_ (0) =0.15. We show two staring values for the mean phenotypic trait: 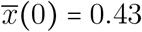 and 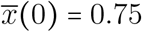. 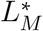 represents the mutation load at equilibrium, which is the difference between *β*_0_ and the equilibrium mean transmission rate 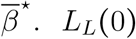 and *L*_*M*_ (0) respectively show the initial lag load and mutation load for the scenario 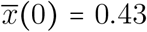.The parameters were *b* =3, *d* =1, *β*_0_= 4, *α* =1, *s*_*β*_= 1, *U*= 2, *f*= 0.4, *λ*= 0.05, *n*= 5. Initial conditions were *I*(0) = 0.1, *S*(0) = *b*/*d*

The transient dynamics is generally well described by (8) but, when selection *s*_*β*_ is strong and/or the effect of mutation *λ* is large, the phenotypic distribution does not remain Gaussian during adaptation. We show in the Supplementary Information (see equations (S38) and (S39)) that *K*_3_ and *K*_4_ are expected to build up during the transient phase of adaptation. Crucially, these higher moments increase the phenotypic variance transiently, which speeds up the adaptation of the virus population (see **Figure S1**). In the long-term, however, we expect *K*_3_ → 0 and 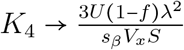.Consequently, when these higher moments of the distribution are accounted for, the phenotypic variance is lower at the endemic equilibrium (**Figure S2**). Hence, as pointed out in the previous section, the critical mutation rate is higher when *K*_4_ is accounted for (compare equations (12) and (15)).

Yet the epidemiological dynamics of the virus population is driven by 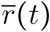 and not by 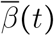.To better understand the dynamics of viral adaptation, it is useful to decompose the dynamics of viral mean fitness *r t* into separate effects following the framework of Gandon & Day [39] (**Figure S4**):

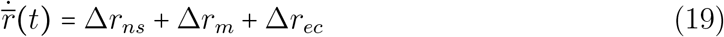

where Δ*r*_*ns*_, Δ*r*_*m*_ and Δ*r*_*ec*_ refer to the changes in mean fitness due to natural selection, mutation and environmental change, respectively. In our model that these different components of the dynamics of adaptation can also be expressed as:

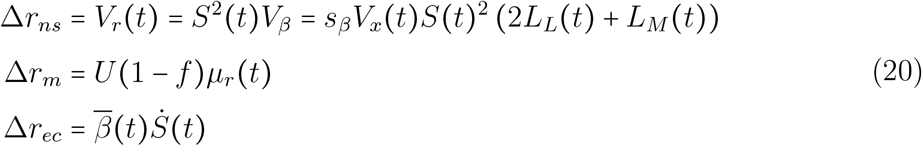

First, as expected from Fisher’s fundamental theorem, the change of mean fitness from natural selection is always positive and equal to the variance in fitness *V*_*r*_(*t*)

(**Figure S4**). This variance increases with the lag load *L*_*L*_ (*t*) (the farther a phenotype is from the optimum, the larger the strength of selection towards this optimum) and the mutation load *L*_*M*_ (*t*) (even if this load has a negative impact on mean fitness, it has a positive influence on the speed of adaptation). Selection is also fueled by the population of susceptible cells and the phenotypic variance, scaled by the shape of the fitness landscape, *s*_*β*_.

Second, the effect of mutations on mean fitness is simply equal to the influx of non-lethal mutation *U(*1 − *f)* multiplied by the mean effect of mutations on fitness *µ*_*r*_ (recall that *µ*_*r*_ (*t*) *= µ*_*β*_*S* (*t*)). Because *µ*_*r*_ <0 (see equation (7)), the direct effect of the mutation process is always negative on the dynamics of adaptation in our model (**Figure S4**). Note that this quantity is exactly equal to the drop in mean fitness in mutation accumulation experiments where the radical bottlenecking at each passage ensures that natural selection does not operate (because the variance in fitness *V*_*r=*_ 0). As expected, lethal mutagenesis occurs when the positive effect of natural selection is overwhelmed by the negative effect of deleterious mutation [12, 26]. But lethal mutagenesis may also be driven by a degradation of the within-host environment of the pathogen which we discuss next.

The third term of equation (2.3) accounts for the environmental change consecutive to a drop in the density of susceptible cells. This final term can be either positive or negative, depending on the change in the density of susceptible host cells (**Figure S4**). During the initial phase of an infection, the density of susceptible cells is expected to drop and to have a negative impact on the growth rate of the epidemic (density-dependent regulation). In contrast, during the initial phase of therapy, drugs are expected to reduce the density of infected cells and, consequently, the density of susceptible cells may increase. We illustrate below how this within-host epidemiological feedback may affect the feasibility of lethal mutagenesis to treat viral infections.

### 2.4 Within-host dynamics during and after a mutagenic treatment

A drug may act in, at least, two different ways in our model. First, a drug may have a mutagenic effect and act via an increase of the mutation load *L*_*M*_ of the virus population. Second, a drug may move the fitness optimum away from the virus population and increase the phenotypic distance 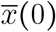. In the following we explore how these two effects act on the transient within-host dynamics of the virus in the WSSM regime. For simplicity, we assume that the other parameters of the model are unaffected by the use of the drug.

**Figure 5** shows virus dynamics during and after the start of drug therapy using the approximation which accounts for the moments *K*_3_ and *K*_4_ (see equations (S36)-(S39)). When, the mutation rate induced by the drug is sufficiently high above *U*_*c*_, the drug can drive the virus population to extinction before the end of treatment. But when the mutagenic effect is lower, the initial stress level *x (*0) is low and/or the duration of the treatment is short, the drug may fail to clear the infection and could even lead to higher rates of viral replication and higher viral loads after the end of the treatment. This rebound is due to the increased rate of viral adaptation induced by the mutagen and to the increase of the density of the susceptible host cells during treatment. It would thus be particularly important to monitor the patients before deciding to end the treatment to avoid the risk of recrudescence of the infection if it was not cleared. The risk of viral evolutionary rescue is less likely if we account for the build up of an effective immune response of the host against the virus as in the case of an acute infection where the death rate of the infected cells will increase after some time. For instance, we show in **Figure S5** how host immunity may reduce the risk of recrudescence. In this scenario, a mutagen may be used as a way to buy some time and reduce the viral load before the immune response kicks in and effectively controls the infection.

**Figure 5:**
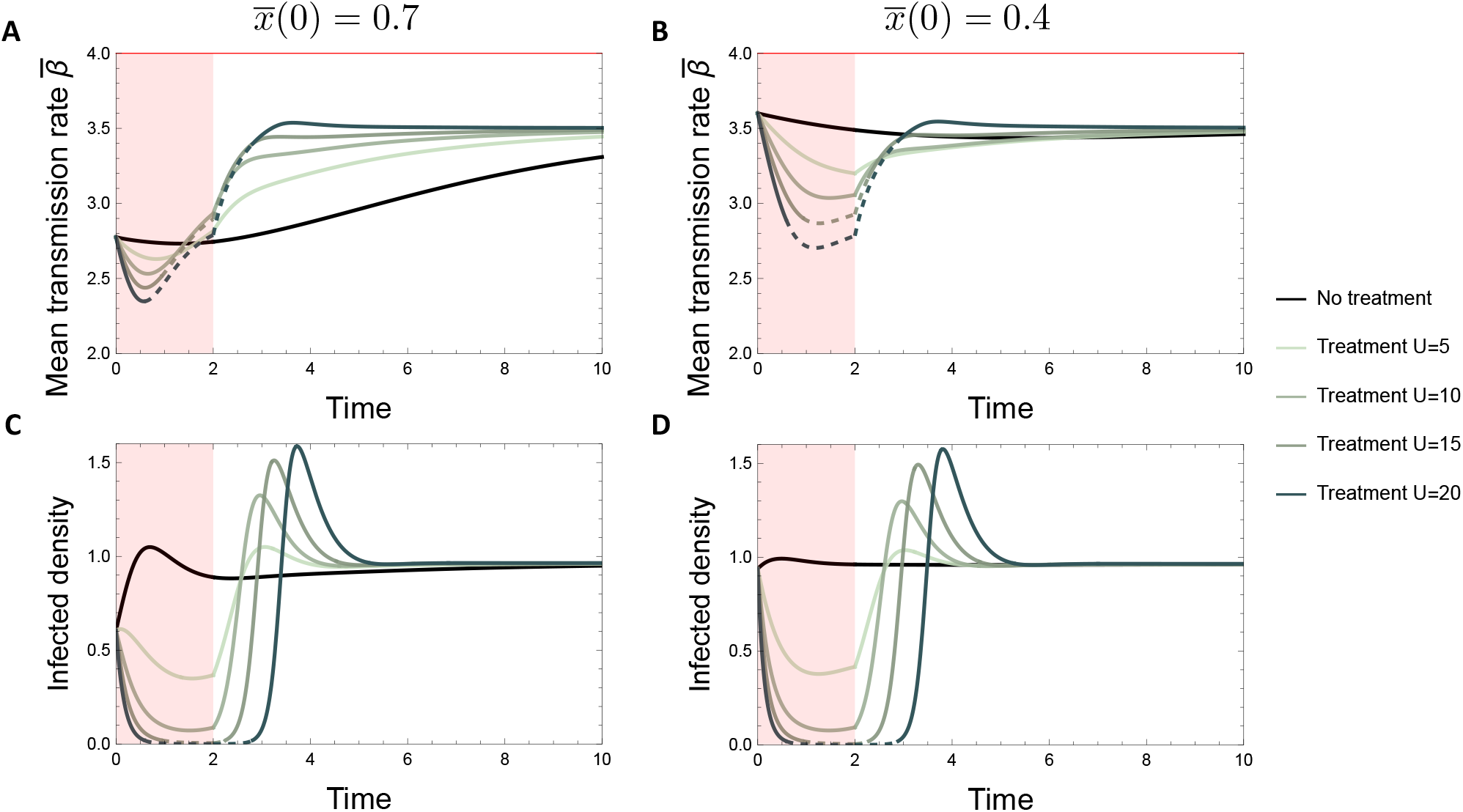
Effect of a mutagenic treatment on chronic infections. The mean transmission rate (A,B) and infected density (C,D) are shown through time depending on an initial distance to the optimum of the pathogen of 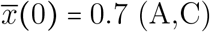 or 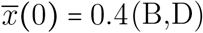 infection, for the WSSM+cumulant model. Simulations are initialized with clones (null variance) and the densities of susceptible and infected cells are initialized with the equilibrium values obtained with *U=* 1. During the treatment (red shaded window), the mutation rate is increased from 1 to the value of *U* specified on the right of the plots. The computed deterministic critical mutation rate is *U*_*c*_ 17.4. The treatment stops at time *t=* 3 where the mutation rate goes back to 1. When the Infected density drops below an arbitrary value of *ϵ=*0.02, the corresponding section of the Infected density and mean transmission rate curves become dashed. This highlights how the changes in mean transmission rate observed in these time frames are highly dependent on escaping stochastic extinction when density is low. The horizontal red line in (A,B) shows the maximum transmission rate *β*_0_. The parameters used were: *b*= 3, *d* =1, *β*_0_ =4, *α* = 1, *f* = 0.4, *λ* = 0.05, *n* = 5.

The above numerical exploration of the effect of a mutagenic drug on within-host viral dynamics ignored the influence of demographic stochasticity. Demographic stochasticity is expected to be particularly high when the viral load drops to very low levels. We explore the influence of demographic stochasticity in **Figure 6** by computing the probability of evolutionary rescue after the start of the mutagenic treatment as a function of the viral mutation rate *U* induced by the drug and *x* 0 the initial stress. Note that we use higher values of x(0) compared to previous figures to obtain cases where initial fitness is negative and thus observe potential evolutionary rescue. As expected, viral extinction occurs when *U* >*U*_*c*_ and larger values of *x (*0) always promote viral extinction. Note, however, that when *x (*0) is sufficiently large, the virus population may only survive when the mutation rate is above a minimal mutation rate *U*_*min*_. Indeed, intermediate mutation rates (i.e. *U*_*min*_ <*U*< *U*_*c*_) can speed up viral adaptation and allow the viral population to avoid extinction at the beginning of the treatment without inducing an unbearable mutation load at equilibrium. The influence of the mutation rate *U* on the probability of extinction has been analysed in [40] without a demographic feedback. Our simulation models accounts for the within-host epidemiological dynamics and show that larger fraction of lethal mutants can have a massive impact on the interaction between the effect of *U* and the initial stress level *x*(0). As expected, the effects of demographic stochasticity are amplified by smaller host and pathogen population size (**see Figure S6**). To appreciate the effects of the environmental feedback of the density if susceptible cells, we show in **Figure S7** scenarios of evolutionary rescue with the same parameters but a constant susceptible density of *b/ d* that show increased probabilities of rescue overall.

**Figure 6.**
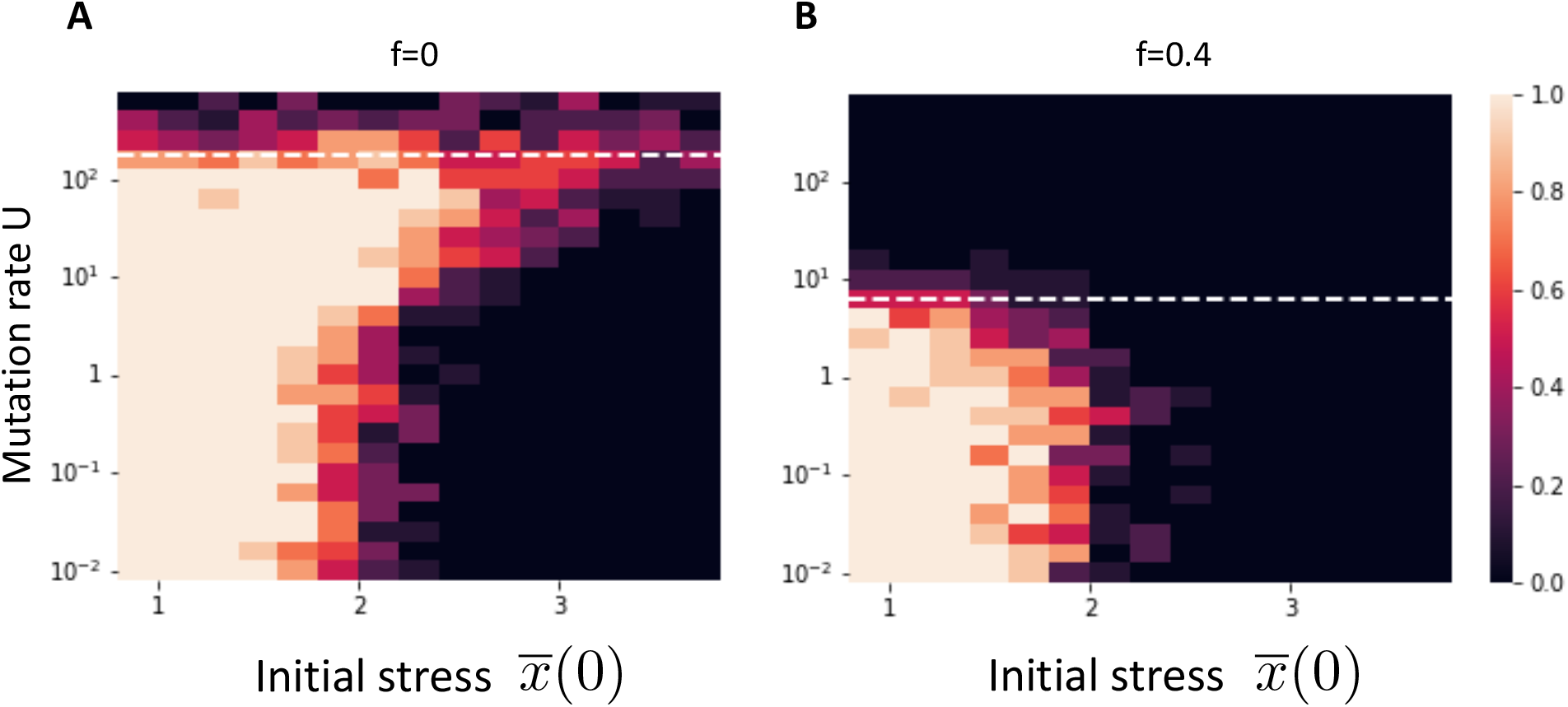
Probability of evolutionary rescue with (A) and without (B) lethal mutations. The simulations are initialized with the equilibrium values of *S,I* and variance *σ* for a mutation rate of *U =*10 ^−2^. The color scale represents the proportion of simulations in which the infected populations survived. There are 10 simulations per parameter combination. The horizontal dashed white line represents the value of critical mutation rate *U*_*c*_ above which the infected population goes to extinction in the deterministic model. The parameters used were: *b* =2, *d* =1, *β*_0_= 4, *α*= 4, *λ*= 0.05, *n* =2, *s*_*β*_ =0.5, 𝒜 1000, *τ* = 0.05, *t*_*max*_ = 50. Note that we take *n* = 2 to reduce simulation time.

## Discussion

To explore the validity of the lethal mutagenesis hypothesis we need to understand the within-host dynamics of a viral population exposed to a high rate of mutation. We developed a model to track both the epidemiological and evolutionary dynamics of the virus. Viral adaptation is assumed to take place in a fitness landscape with a single fitness optimum. Mutations allow the virus phenotype to move in this landscape on *n* independent dimensions. Each dimension is under stabilizing selection and contributes to the within-host viral transmission rate *β*_x_. We couple this model of adaptation with an epidemiological model of within-host dynamics, in which the availability of susceptible host cells feeds back on the growth rate of the virus. We model the dynamics of the phenotypic distribution through the derivation of the cumulants of the distribution [35]. Our model accounts for the occurrence of both lethal and non-lethal mutations as observed in empirical measures of distribution of fitness effects in viruses [4, 11]. Lethal mutations have a strong but instantaneous effect and they can be viewed as an extra source of viral mortality in our model. Non-lethal deleterious mutations, in contrast, can accumulate over time and bring down the mean transmission rate of the virus.

### Critical mutation rate

Our model can be used to predict the long-term evolutionary equilibrium of the virus population. In particular, we can predict the critical mutation rate *U*_*c*_ above which the mutation load is so high that virus population is driven to extinction. We derive approximations for *U*_*c*_ under different regimes of selection. First, when the effect of mutation *λ* is small and selection *s*_*β*_ is relatively weak relative to the mutation rate (WSSM regime), the equilibrium mutation load *L*_*M*_ (combine equations (9) and (11)) is driven by the effects of non-lethal mutations which depends on the number *n* of phenotypic dimensions (i.e. complexity). Indeed when phenotypic complexity is high, the proportion of beneficial mutations decreases in favor of deleterious mutations. The mutation load depends also on *λs*_*β*_ which measures the product of the phenotypic effects of mutations *λ* and the strength *s*_*β*_ of stabilizing selection.

As expected, larger values of *λs*_*β*_ reduce *U*_*c*_ (**Figure S3**). Second, when selection is strong relative to the mutation rate, the phenotypic variance of the viral population is lower than in the WSSM regime and we can use other approximations to predict the critical mutation rate. We showed that we can account for the fourth cumulant of the phenotypic distribution to improve our predictions but, when selection is much stronger (condition (17)), we can use the classical “House of Cards” (HC) approximation. In this regime, the effects of non-lethal mutations is so strong that they can be viewed as effectively lethal in the WSSM regime (compare equations (13) and (18)).

Crucially, all the parameters required to predict the critical mutation rates of a virus can be obtained experimentally (see **Supplementary Information S2.9**). In **Table 1** we combine different experimental results to derive the critical mutation rates *U*_*c*_ for six different viruses. Of course, our predictions should be treated with caution because they are based on a simple within-host model of transmission and we ignore the influence of host immune system. Yet, its is interesting to see that some convergent results emerge from the analysis of this very different viruses. First, we find that the critical mutation rate is always higher for the WSSM than for the HC approximation. This indicates that, for all these viruses, the WSSM approximation is more relevant to predict *U*_*c*_ (see **Figure 3**). Second, if we compare this critical mutation rate with the natural mutation rate of these viruses we find that lethal mutagenesis requires a 14-to 3080-fold increase of the genomic mutation rate. Yet, available estimates of the mutagenic effect of nucleoside analogues like ribavirin, favipiravir and molnuparivir show that they increase viral mutation rate by a factor of 10 at most [41–43]. This suggests that the increased mutation rate triggered by the currently available mutagenic drugs is too low to eradicate a viral infection by lethal mutagenesis only (i.e., via an *indirect effect* where the increased rate of mutation reduces, 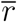, the mean growth rate of the virus population). Yet accounting for the additional *direct effects* of the drugs and/or the host immune response on, *r*_0_, the maximal growth rate of the virus are expected to reduce dramatically the critical mutation rate needed to eradicate the virus population. The above discussion of the feasibility of lethal mutagenesis, however, is based on the long-term evolutionary equilibrium and on the principle that viral eradication is expected to occur when *U> U*_*c*_. But the time needed to eradicate the virus population could be longer than the duration of the treatment. Hence, in practice, viral recrudescence may occur after the end of a treatment with a strong mutagen (**Figure 5**). Besides, we also showed scenarios where the virus population can go extinct even when *U*< *U*_*c*_ (**Figure 6**). In other words, the condition *U> U*_*c*_ is neither necessary nor sufficient to eradicate the virus. To increase our ability to predict the viability of the virus population we need to understand the transient within-host dynamics of the virus.

### Transient within-host dynamics

We studied the transient within-host dynamics of the virus in the WSSM regime where the phenotypic distribution remains Gaussian during the adaptation of the virus population. When selection is stronger, however, the distribution can move away from the Gaussian distribution (equations (S36)-(S39)) [35, 44–46]. The transient build up of Skewness increases the variance in fitness and tends to speed up the rate of adaptation (**Figure S1**). The analysis of the transient dynamics under the HC regime (when selection is very strong) remains to be carried out.

This framework allowed us to explore the effect of a mutagenic drug on viral dynamics. The drug is expected to affect the epidemiological dynamics (both the density of infected and uninfected cells) and the evolutionary potential of the virus. We showed that if the treatment is stopped before the eradication of the virus, the infection could rebound and reach very high viral loads (even if the drug is an effective mutagen and *U> U*_*c*_). The risk of this viral rebound, however, is limited in hosts that are able to mount an effective immune response (see **Figure S5**)). Yet, the risk of rebound could be high for immunodepressed hosts and chronic infections. Hence, it is particularly important to monitor the patients treated with a mutagen after the end of the treatment.

When we explored the robustness of our predictions in a finite population of host cells we found that the virus could go extinct even when *U< U*_*c*_. These extinctions occur when the viral population is initially far away from its fitness optimum. The lower the initial level of adaptation, the more likely the virus population may be driven to extinction by the drug. This could occur if a mutagenic treatment is combined with a direct reduction in fitness by another drug. This effect is particularly strong when the fraction *f* of lethal mutations is high because the evolutionary potential is limited in this case (**Figure 6**).

### Concluding remarks

In our model, mutations can only affect the within-host transmission rate of the virus. However, other life-history traits could be involved in viral fitness and it is necessary to describe the joint distribution of mutation effects on these different life-history traits. For instance, we could study the joint evolution of transmission *β* and virulence *α*. Interestingly, the effect of transmission on fitness is scaled by the density of infected cells, but not virulence (see equation (5)). We may thus expect epidemiological feedbacks to have different effects on the evolution of transmission and virulence [39, 47]. Crucially, the optimum phenotype may differ when selection acts on transmission or on virulence which may lead to a trade-off between the two traits (i.e. impossible to maximize transmission and minimize virulence). It will be interesting to test the robustness lethal mutagenesis when multiple life-history traits are jointly evolving, but we leave this for future studies.

Our model focuses on within-host dynamics but if a patient treated with a mutagenic drug does not fully clear the infection, drug-mutated viruses can be transmitted to a new hosts. For example, the use of molnupiravir, an antiviral mutagen used against SARS-CoV-2 in some countries is associated with the rise of specific G-to-A mutations in the virus in these countries [42]. In principle, some of these mutations may affect the between-host fitness of the virus (e.g. higher between-host transmission, ability to evade host immunity and infect vaccinated hosts). These mutations may not necessarily affect within-host dynamics but they could have dramatic consequences on the emergence of new variants. Hence, a more comprehensive measure of evolutionary safety of lethal mutagenesis therapy should also consider the cumulative number of mutants that may be transmitted to new hosts [48]. An in depth analysis of the within-host and between-host consequences of mutagenic drugs remains to be explored.

## Acknowledgements

We thank Guillaume Martin and Ophélie Ronce for numerous discussions and comments on earlier versions of this work.

## Supplementary Information

In this supplementary information we detail the derivation of the theoretical results presented in the main text. In section 1, we start by presenting the main assumptions on the fitness landscape, the effects of mutations and the epidemiological dynamics. In sections 2 and 3, we present different ways to analyse the joint epidemiological and evolutionary analysis of this model and obtain analytical approximations. In section 4, we detail the numerical model used to check our analytical approximations.

### S1. The model

We model the within-host dynamics of of a viral population that spreads in a large population of susceptible cells. We assume that different strains of the virus may have different phenotypes. Each phenotype **x** is defined as a vector of size *n*, representing *n* independent continuous phenotypic traits. Each host cell is assumed to be infected by a single strain (no multiple infections) and the total density of infected cells is noted *I =*∫*I*_**x**_*d*^*n*^**x** and *p*_x_=*I*_x_/ *I* is the frequency of cells infected by strains with phenotype **x**. We assume that the viral transmission rate *β*_x_ is governed by the values of the *n* underlying phenotypic traits through a Gaussian transmission function. This Gaussian is well approximated with a quadratic function when phenotypes are not too far from the optimum, ensuring no negative transmission rate values. This quadratic form yields:

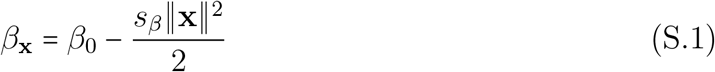

where ∥**x**∥ is the norm of vector **x** and *s*_*β*_ measures the strength of selection towards a single optimum at **x** = (0, 0, …, 0) of maximum transmission rate *β*_0_ (see **Figure 1** in the main text). Different virus strains may have different transmission rates *β*_**x**_ and the average transmission rate is given by:

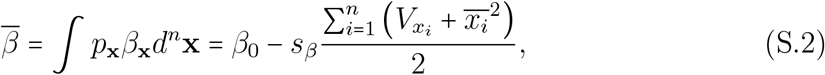

where 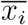 and 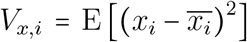 are respectively the mean and the variance for the phenotypic trait *i*.

We model the within-host spread of the viral population through the dynamics of *S* and *I* the densities of uninfected and infected cells, respectively. We assume there is a constant influx *b* of susceptible cells which die at constant rate *d*. A cell infected with a virus strain with phenotype **x** infects new susceptible cells at a rate *β*_**x**_ and dies at a rate *d+ α* where *α* measures the increased mortality rate induced by the infection. Viral mutations may occur at a constant rate *U* and among those mutations a fraction *f* may be lethal for the virus. Consequently, the rate of lethal mutation *Uf* may be treated as an additional mortality term for infected cells. Note that, in the following, the dependence to *t* of most dynamical variables is dropped to simplify the notation. This yields the following dynamical system:

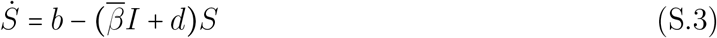

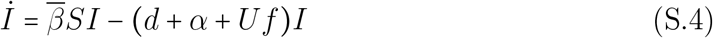

In the absence of the pathogen the density of the uninfected cells is equal to *S*_0_ =*b/ d*. In such a naive host population a pathogen with average transmission rate 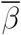 can spread if and only if its basic reproductive ratio:

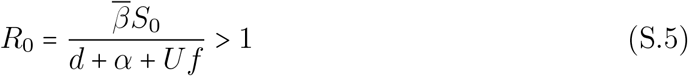

In other words a condition for the viability of the pathogen population at equilibrium is 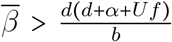.We can write the dynamics of *I*_x_ the density host cells infected by strain *x* using the following integro-differential equation:

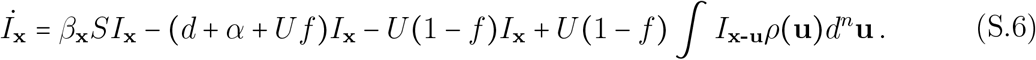

The first two terms describe the *birth rate* and the *death rate* of infections, respectively. The following term describe the mutation away from strain **x**, and the last term describes the mutations from all other strains to strain **x**. At rate *U* (1 −*f*) (the effect lethal mutations have already been accounted in the *death rate* of infections), strain **x-u** mutates to phenotype **x** with a probability *ρ (***u**) following an isotropic multivariate normal distribution with mean zero and variance *λ. λ* thus refers to the variance of the phenotypic effect on each phenotypic dimension.

### S2. Dynamics of the cumulants of the distribution of phenotypes

In this section we use the integro-differential equation (S.6) to derive the cumulants of the distribution of phenotypes **x**. Cumulants are quantities directly related to the moments of the distribution as we show below. We use cumulants over moments because of a property of Gaussian distributions: their cumulants of order >3 are equal to zero. We will use this property for “moment closure”, ie. to neglect higher order cumulant in our dynamical equations to obtain a closed system of differential equations. Interestingly, one can relax this Gaussian approximation to a certain degree by taking into account these high order cumulants. In all our derivations we will assume that cumulants of order ≥5 can be neglected.

In the following, 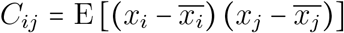 denotes the phenotypic covariance between traits *i* and *j*, while 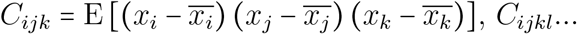 denote higher-order moments. Cumulants *K*_*U*_ can be defined in terms of moments *C*_*U*_ from the definition of the cumulant generating function. In particular, we have (see also [1, 2]) :

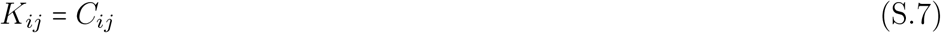

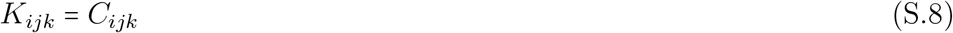

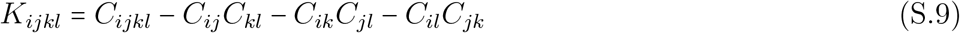

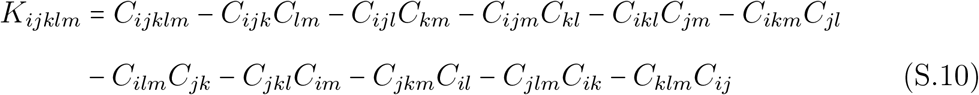

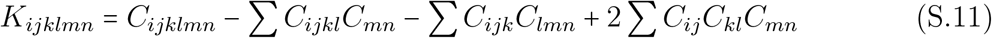

where in the last expression the sums include all terms obtained by permuting indices. If the distribution of phenotypes is Gaussian, all cumulants of order ≥ 3 are zero. In the following, we derive expressions for the dynamics of mean phenotypes 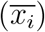,phenotypic variances and covariances (*C*_*ij*_) and cumulants of order 3 and 4 (*K*_*ijk*_, *K*_*ijkl*_).

Changes in phenotype frequencies *p*_x_ = *I*_x_ (*t*)/ *I* (*t*) among the non-lethal viruses are given by:

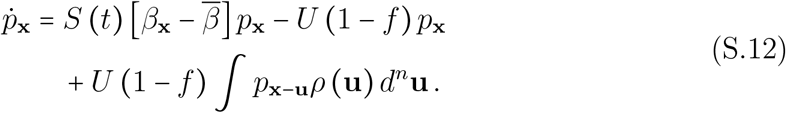

#### S2.1. Dynamics of mean phenotypes

Changes in mean phenotypes are given by:

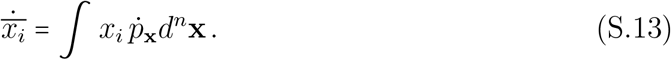

From equations S.12 and S.2, this gives:

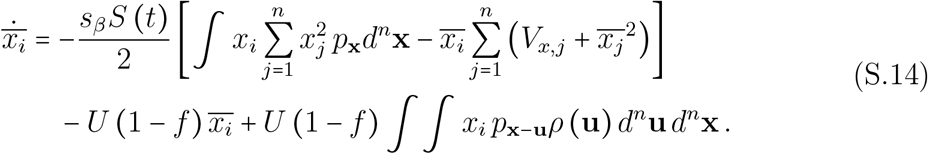

The last line of equation S.14 cancels, while the integral on the first line may be written as:

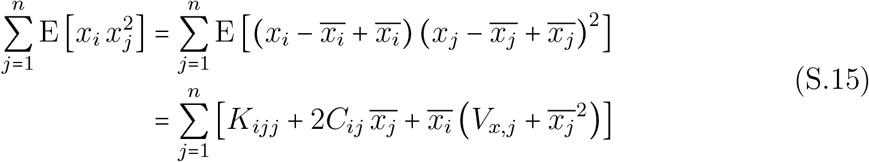

Finally giving:

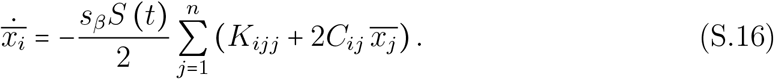

#### S2.2. Dynamics of second moments

Similarly, changes in phenotypic variances and covariances are given by:

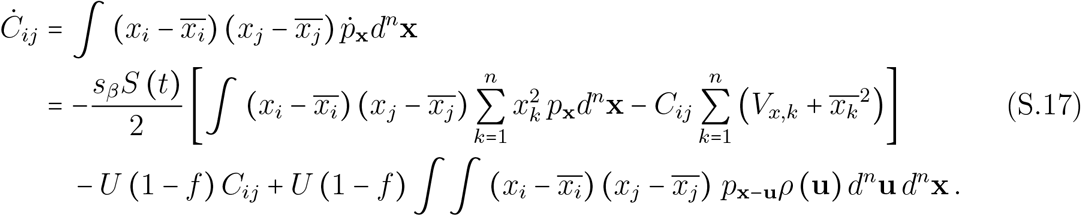

The integral in brackets on the second line of equation S.17 may be written as:

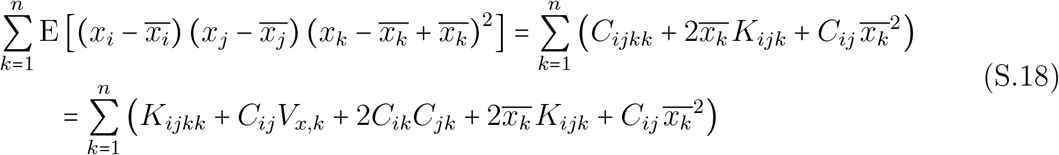

(using equation S.9) while the double integral on the third line of equation S.17 may be written as:

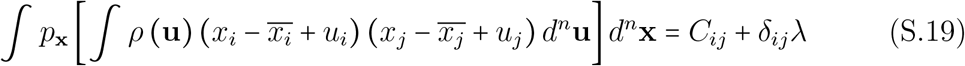

where *δ*_*ij*_ = 1 if *i* = *j*, and 0 otherwise. Putting everything together yields:

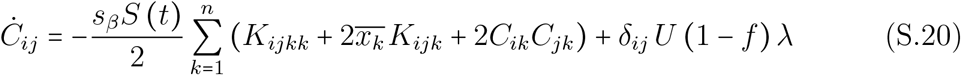

#### S2.3. Dynamics of third cumulants

The change in *K*_*ijk*_ = *C*_*ijk*_ is given by:

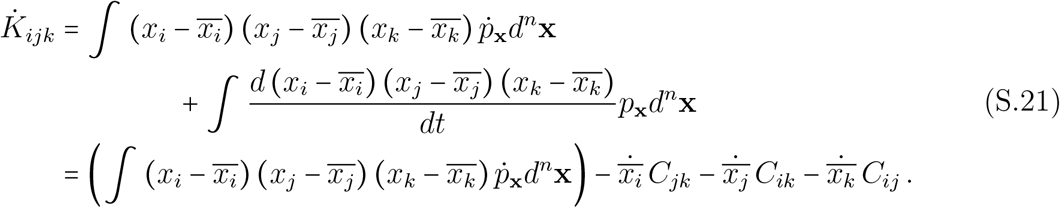

As before, the integral on equation S.21 writes:

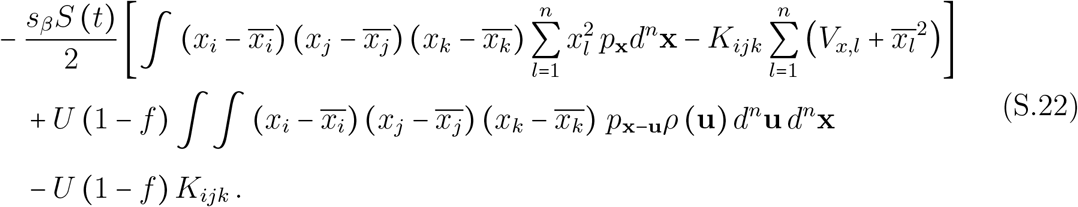

The last two lines of equation S.22 vanish under the assumption that third moments of the distribution of mutational effects *ρ* are zero, while the integral on the first line can be written as:

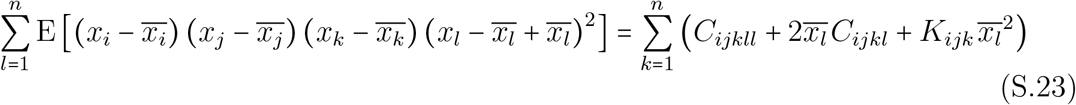

The moments *C*_*ijkll*_ and *C*_*ijkl*_ can be expressed in terms of second-order moments and third / fourth-order cumulants using equations S.9 and S.10 (assuming that the fifth-order cumulant *K*_*ijkll*_ equals zero). Putting everything together, one finally obtains:

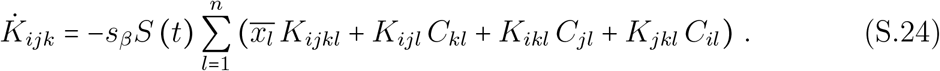

#### S2.4. Dynamics of fourth cumulants

Finally, from equation S.9 we have:

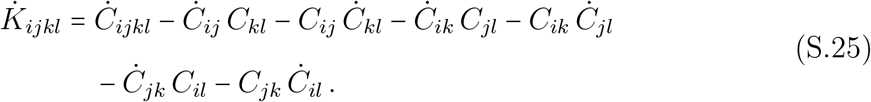

The change in *C*_*ijkl*_ is given by:

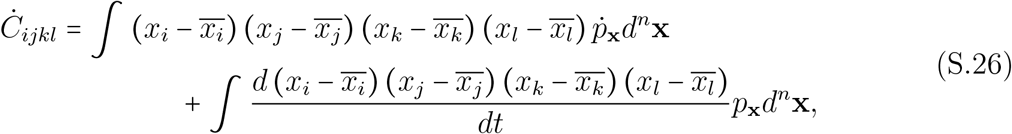

the second line of equation S.26 being equal to:

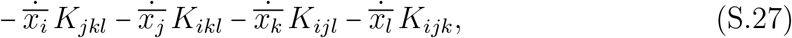

while the first line is:

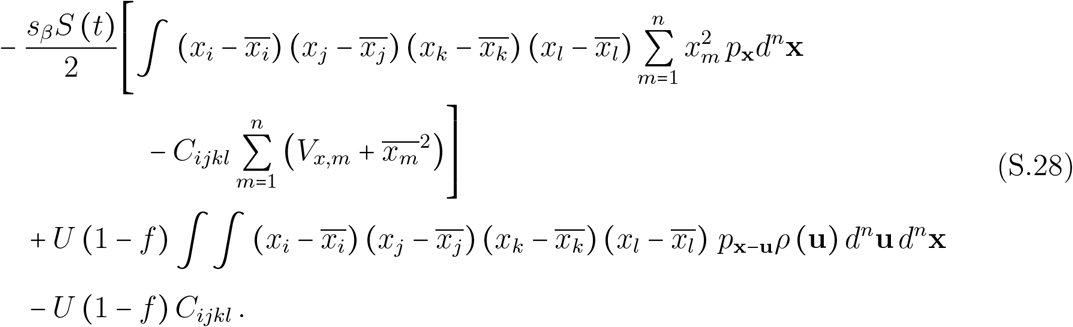

The first integral is computed as before, and yields:

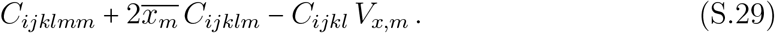

The moments *C*_*ijklmm*_, *C*_*ijklm*_ and *C*_*ijkl*_ can be expressed in terms of second-order moments and third / fourth-order cumulants using equations S.9, S.10 and S.11 (assuming *K*_*ijklm*_ = 0, *K*_*ijklmm*_ = 0). From this, one obtains that the change in *K*_*ijkl*_ is given by:

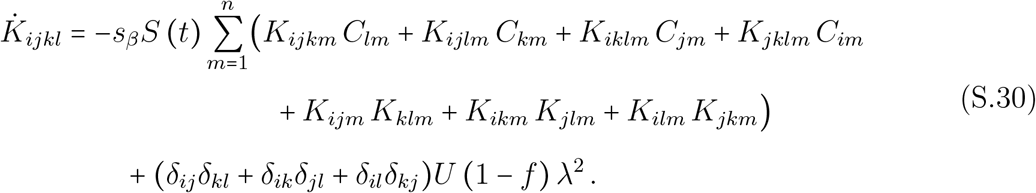

#### S2.5. Simplification for isotropic mutation and selection

If the initial distribution of phenotypes **x** is a multivariate normal distribution with mean 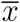 and variance *V*_*x*_ in all dimensions, we have initially independence of all phenotypic trait *x*_*i*_. This implies that only cumulants of the same phenotypic trait (eg. *C*_*iiii*_) are non-zero. If these cumulants are initially zero, one can check with their dynamical equations that they will remain zero throughout the dynamics because selection and mutation do not generate anisotropy. Overall, this means that the dynamics of the distribution of phenotypes **x** only depends on the dynamics of *x*_*i*_, *K*_*ii*_, *K*_*iii*_, *K*_*iiii*_. We thus simplify our notations using, for all phenotypic dimensions *i* :

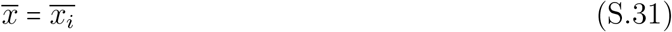

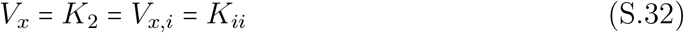

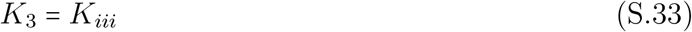

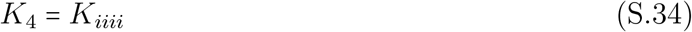

First, the isotropy in phenotypic traits allows us to rewrite the expression of the mean transmission rate as:

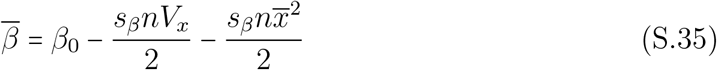

Finally we can write the dynamical system for the cumulants as:

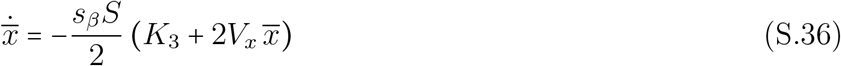

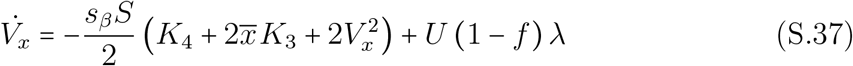

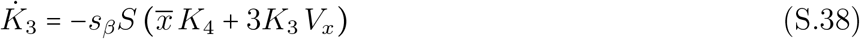

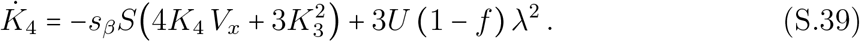

Where *K*_3_ (*t*) and *K*_4_ (*t*) are respectively the third and fourth cumulants of the distribution of phenotypes. Note that this corresponds to the system (7) from the main text for the mean phenotype and the phenotypic variance, but with the additional effect of *K*_3_ and *K*_4_, the third and fourth cumulants of the phenotypic distribution. As expected, when larger effects of mutations can be neglected (i.e. here *λ*^2^ =0), both *K*_3_ and *K*_4_ converge to 0 and we recover system (7) from the main text. Yet, when *λ*^2^ cannot be neglected, the influx of new mutations increases the cumulant *K*_4_. A positive *K*_4_ means that the phenotypic distribution is both more peaked around the mean phenotype and more heavily tailed, with less intermediate phenotypes than in a Gaussian distribution. This fourth cumulant is also expected to generate transiently some skewness *K*_3_ (which sign is inverse to the sign of 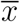). This is expected to transiently increase the variance in fitness and speed up viral adaptation. In the long run, however, the skewness *K*_3_ is expected to vanish as the viral population gets closer to the optimal phenotype. Yet, the kurtosis remains even at the equilibrium and it decreases the equilibrium phenotypic variance. This lower phenotypic variance results in a lower mutation load. As discussed above with the effect of lethal mutations, increasing *λ* has a similar effect as in the WSSM because it increases the efficacy of natural selection and allows the purging of deleterious mutations.

#### S2.6. Equilibrium phenotypic variance

The equilibrium values of 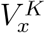 (the superscript *K* for the cumulant approach) and *K*_4_ must satisfy:

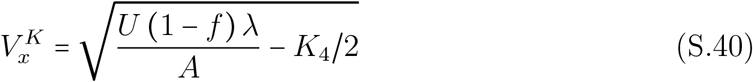

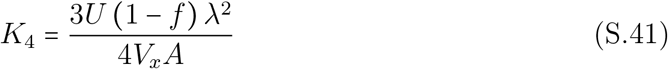

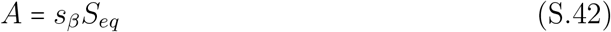

We can express the equilibrium density of susceptible cells as a function of the equilibrium phenotypic variance (using (4) and (35) when the pathogen has reached an evolutionary and epidemiological equilibrium), yielding:

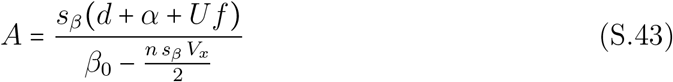

If we neglect *λ*^2^ terms (WSSM regime) we have *K*_4_ =0 and we get the following expression for equilibrium variance:

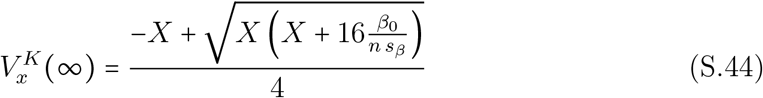

with:

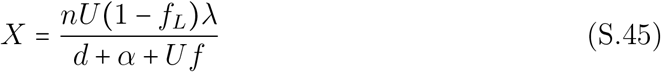

If we want to account for *λ*^2^ terms, we have to consider higher order mutational effects directly with equation (S.40). We failed to obtain a simple expression for the equilibrium variance in this case, but we can study the predictions numerically (Figure S2 in Main text). Interestingly, when *λ* or *s*_*β*_ are very high, the above dynamical system does not reach a stable equilibrium with a positive *V*_*x*_. With (40) and (41) we obtain the following necessary condition for a positive *V*_*x*_ to exist: 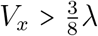.But we can narrow down the condition of validity of this approximation after noticing that only the upper part of the solution of *V*_*x*_ as a function of *s*_*β*_ is valid (see figure S2). Finding the limit value of *V*_*x*_ yields a new condition of validity: 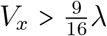.In other words, the standing phenotypic variance has to be substantially higher than the mutational variance. This also yields a threshold value for *s*_*β*_:

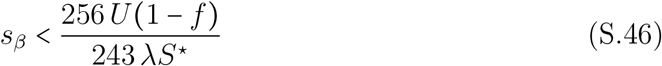

Note how this value is not directly dependent on the phenotypic complexity *n* (except indirectly through *S*^∗^), contrary to the threshold value for the transition from the WSSM to the HC approximation (main text equation (15)). With higher values of *n*, the range of validity of this approximation will increasingly overlap with the HC validity range (**Figure S3**).

#### S2.7. The House of cards model

On the other side of the spectrum, there are regimes where mutations are rare but with larger effects. For this case, the ‘House of Cards’ approximation has been developed by Turelli [3], based on the assumption that the effect of a mutation is independent on the original phenotype in which it appeared. In this case equation (S.12) becomes :

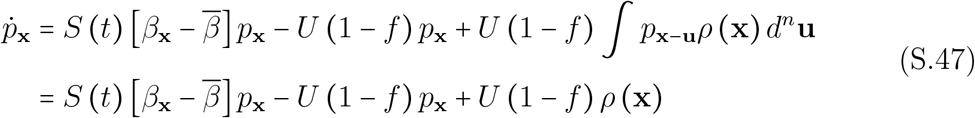

At equilibrium, this leads to :

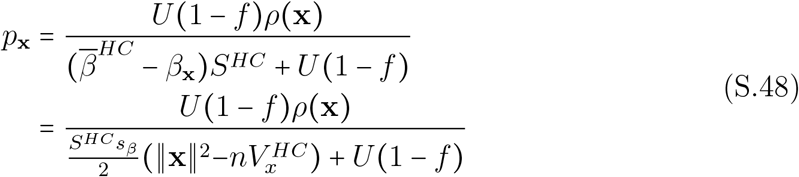

With *p* being a probability density function, we get that

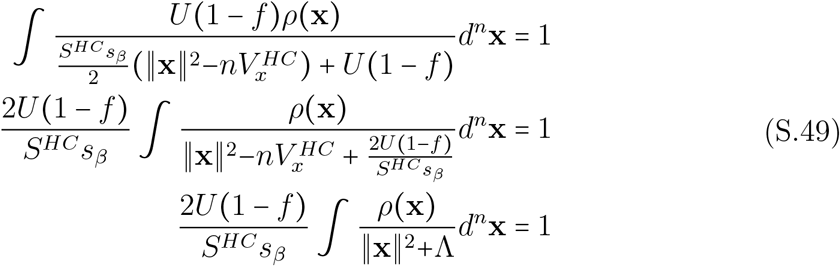

with 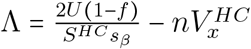.Noting in equation (S.48) that the right-hand ratio must be positive and that *U* (1 − *f*)*ρ*(*x*) > 0 we get in particular with *x* = 0 that

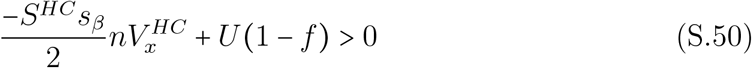

from which follows an upper bound of the equilibrium variance

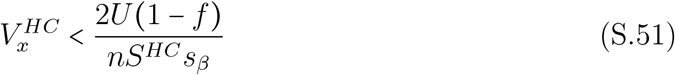

The main difference with our previous results is that the equilibrium variance is independent of the mutational variance *λ*. We use our simulation model to compare the different expressions for equilibrium variance in the different regimes. We see that the cumulant approach up to *K*_4_ provides the best fit when *s*_*β*_ is high (i.e., when selection is strong). In case of extremely low selection, this approximation becomes equivalent to the WSSM regime because the term in *λ*^2^ vanishes. However, the fourth cumulant approximation crumbles with higher selection where it yields negative values for the variance. In this case, the House of Cards approximation yields the best result. Indeed in this regime, the distribution of phenotypes is more peaked around the optimum and the distribution is very far from Gaussian, which is the basis of the WSSM approximation. The condition for the switch between a WSSM regime and the House of Cards regime is given by setting 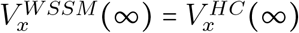 and solving for the mutation rate *U*, which yields:

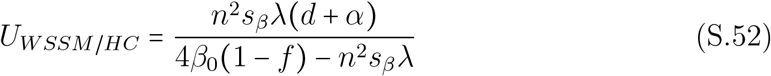

#### S2.8. Critical Mutation Rates

The critical mutation rate is the value above which the pathogen population goes extinct because the mutation load is too high (i.e., the transmission rate is too low). In this case the equilibrium density of susceptible cells is:

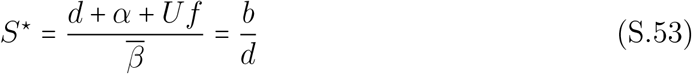

and using (S.35) at an evolutionary equilibrium (i.e., *x* = 0):

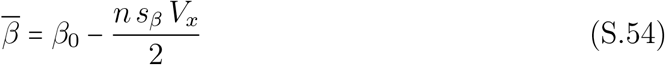

The critical value of the phenotypic variance is thus:

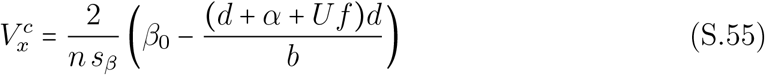

Then one can use (S.44) to derive the critical value of the mutation rate *U*_*c*_. We find that when *f* > 0:

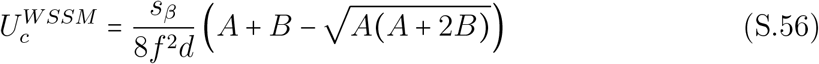

with:

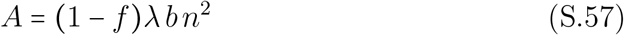

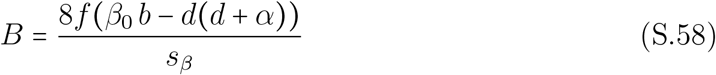

In the special case when *f* = 0:

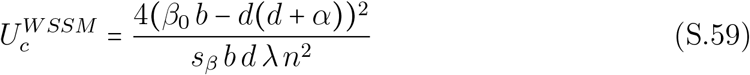

Similarly, we can compute the critical mutation rate In the House of Cards regime which gives:

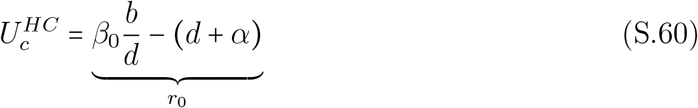

Strinkingly, this threshold of critical mutation rate is not dependent on other parameters affecting the effect of mutations *λ, s*_*β*_, *n* and *f*. The critical mtuation rate only depends on demographic parameters. This means that in this regime, mutations that we model as ‘non-lethal’ have such strong effects that they are *de facto* lethal when computing a critical mutation rate. In fact, this critical mutation rate is exactly the one predicted in the WSSM regime with equation (S.56) when all mutations are lethal i.e. *f=* 1.

We cannot get an explicit expression for the critical mutation rate in the model where we consider more cumulants. We can however get these values numerically and we compare the three models in the main text.

#### S2.9. Linking empirical data with our predictions

In this section, we detail how available empirical data can be used to predict the critical mutation rate *U*_*c*_ derived in the previous section for different viruses (**Table 1**). First, combining equations (1) and (5) we need to Since empirical we can write:

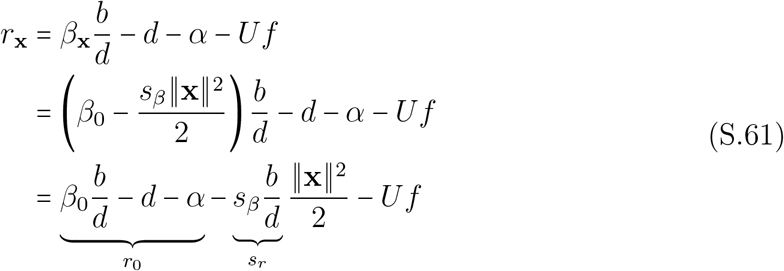

where 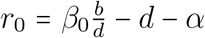 is the maximal growth rate of the virus and 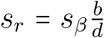 measures the intensity of selection. Note that we compute the fitness when *S =b/ d* because we want to determine *U*_*c*_ (see section S2.8).

In the WSSM regime, we can rewrite (S.56) as a function of *r*_0_ and *s*_*r*_:

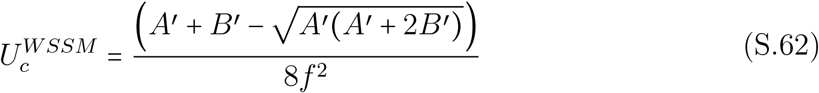

where *A*^′^ = (1 − *f*)*λs*_*r*_ *n*^2^ and *B*^′^ = 8*fr*_0_.

Similarly, in the HC regime, we can rewrite (S.60) as a function of *r*_0_ and *s*_*r*_:

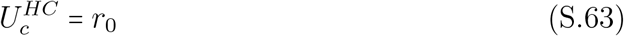

The quantity *r*_0_ can be measured empirically in the absence of a mutagen in a population of otherwise fully susceptible cells. The parameters *f, n*, and the product *λs*_*r*_ can be estimated from the distribution of fitness effects as discussed in Martin & Gandon [4]. In the main text we show that the mean effect of mutations on transmission is *µ*_*β*_ and the mean effect of mutations of non-lethal mutations on growth rate is:

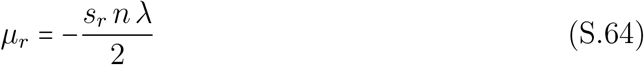

Note that these effects can be rescaled with the maximum growth rate *r*_0_ of the virus which yields:

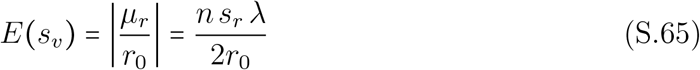

When a phenotype is at the optimum in the FGM, it can be shown that the fitness effect of mutations can be well described with a Gamma distribution. In particular, the shape parameter *α* of this gamma distribution is related to the number of phenotypic dimensions such that *α =n/* 2 [4, 5]. This parameter can be estimated from the data as *α* = 1/*CV* ^2^(*s*_*v*_) where *CV* (*s*) is the coefficient of variation of the fitness effects of non-lethal mutations: 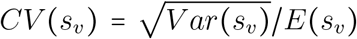.Hence, we can use the observed mean and variance of the fitness effect of non-lethal mutations to obtain:

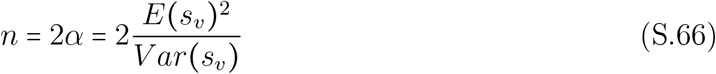

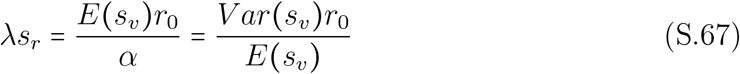

##### Empirical data used in Table 1

We obtained the mutation rates of the five different viruses of **Table 1** from [6]. These rates of mutations are estimated as substitutions per nucleotide and per cell infection (s/n/c). We multiplied these rates by the genome size of each virus to obtain their genomic mutation rate *U*.

The maximal growth rate (*r*_0_), the generation time (*τ*) and the distribution of fitness effects (*E (s*_*v*_), *V (s*_*v*_)and *f*) for Q*β*, Φx174, F1/M13 and VSV were obtained from [7]. These parameters allowed us to obtain *λs*_*r*_ and *n* in **Table 1** using equations (S.66) and (S.67) (see also [4]). The generation time *τ* (a proxy for the duration of cell infection *c* in [6]) was derived using the approximation *τ =*log *R*_0/_ *r*_0_ for virulent phages [4, 7]. Note that F1/M13 are chronic phages (virions are produced without killing the host cell) and the use of the following approximation may be more appropriate *τ =(R*_0_ −1)/ *r*_0_[8] but yields very similar values.

For influenza, the maximal growth rate *r*_0_ was estimated using *in vivo* data from nasal swabs of human infections from [9, 10]. We give the geometric mean of the estimates of *r*_0_ of 6 patients in **Table 1**. We used the expression for *r*_0_ given by [10] and the parameters estimated in [9] with a model that does not account for a delay in virus production. The parameters for the distribution of fitness effects (*E (s*_*v*_*), V (s*_*v*_) and *f*) were obtained from [11]. For the generation time *τ* we give two different estimates. First, we use the approximation *τ= (R*_0_ −1)/ *r*_0_ when the duration of infection is exponentially distributed and second, we use the mean duration of a cell infection *τ* = 1/*δ* because at the beginning of the infection many target cells are available and the cell cycle is mosltly limited by the duration of the cell infection [8]. These two approximations provide a range of values in Table 1. Note that the duration of cell infection used in [6] is 7*h* and falls within this range.

For SARS-CoV-2, we used estimations for the genome size and the mutation rate from [12, 13]. For the distribution of fitness effects, given the remarkably similar values obtained for the five other viruses we took the average values of the above viruses (indicated with a star). The maximal growth rate *r*_0_ was estimated from the doubling times measures in vivo in [14] and the generation time *τ* was estimated in [15].

Next, we use these parameter values in (S.62) and (S.63) to predict the critical genomic mutation under the WSSM or the HC approximation for each virus. Finally, we compute the ratios 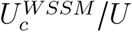 and 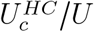 to predict the fold-increase in the rate of genomic mutation required to reach the extinction threshold.

### S3. Deterministic simulations

To check our analytic results, we carried out numerical simulations in two dimensions (i.e., *n=* 2). We build a grid of size *l* ×*l* with *l* an odd number, where each square of the grid corresponds to a phenotype. The phenotypic trait value goes from −*x*_*max*_ to *x*_*max*_ and the phenotype step between each square is 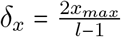 as shown in **Figure S8**. To each square (*i, j)* of the grid is associated at a given time *t* a number of infected individuals *I (i, j, t)* of phenotype (*x*_*i*_, *x*_*j*_).

At each time interval *dt*, susceptible cells *S(t)* grow with a constant rate *b* and die with rate *d*. Infected cells *I(i, j, t)* grow by infecting susceptible cells *S (t)* depending on the transmission rate of their phenotype *β*(*x*_*i*_, *x*_*j*_), and die with *d* + *α* + *Uf*, such that

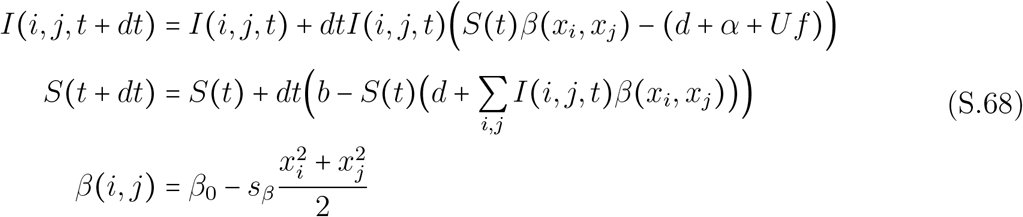

To stick with our analytic model we use a quadratic link function between phenotypes and transmission rate that could lead to negative values of *β*. To add the effect of mutations, we use two different model. We implement the gaussian mutation kernel such that:

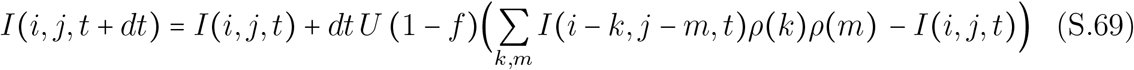

### S4. Stochastic simulations

To explore the effect of demographic stochasticity we carried out stochastic simulations in a finite, but not fixed, host population. The total size of the host population is noted 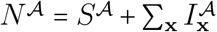,where *S*^A^ refer to the discrete number of susceptible hosts and 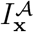 to the discrete number of hosts infected by the pathogen with phenotype **x**. The susceptible hosts immigrate at rate *b*𝒜, where 𝒜is a scaling parameter that indicates the order of magnitude of the arena in which the epidemic occurs ([16, 17]). Hence the total host population varies stochastically in time, but remains of the order of 𝒜. Similarly, we assume that the transmission rate is *β*_**x**_ **/**𝒜so that the basic reproduction ratio is independent of 𝒜. The parameter 𝒜is used to scale the population sizes and thus the impact of stochasticity (lower values of 𝒜increase the amount of demographic stochasticity).

**Table S1** gives a detailed account of the jumps in the Markov chain and their rates. We implement this Markov chain using the *τ* -leap approximation of the Gillespie algorithm ([18]), performing all reactions for an interval of length *τ* before updating the propensity functions. This yields the following procedure:

1. Calculate the event rates *R*_*e*_ (last column of Table S1).
2. For each event *e* generate *K*_*e*_ *=* 𝒫*(R*_*e*_ *τ*), which is a Poisson distributed random variable with mean *R*_*e*_ *τ*. Hence, *K*_*e*_ is the number of times each event *e* occurs during the time interval [*t, t*+ *τ*).
3. Update the state by:

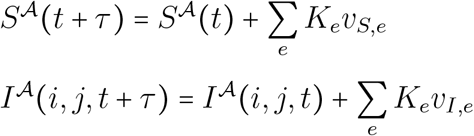

where *v*_*S,e*_ and *v*_*I,e*_ are the change on state variable *S*^A^ and *I*^A^ due to event *e* (second column in Table S1).
4. For simplicity, the dynamics of the susceptible hosts is assumed to be updated in the following way:

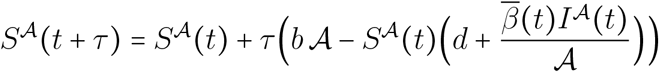

All simulations are initialized at the endemic equilibrium of the deterministic model when *U=* 0.01 and the optimum of the fitness landscape is shifted by 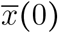,which quantifies the “stress level”. Simulation code is available upon request and will be deposited on github after acceptance of the manuscript. All simulations are done with a maximum time *t*_*max*_ = 50 and *τ* = 0.05. We checked that longer simulation runs (i.e. *t*_*max*_ = 250) had a negligible effect on figure S9.

**Table S1:** We model the epidemic via a Continuous Time Markov Chain (CTMC) with discrete states 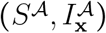.Jumps 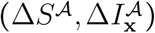 occur at fixed rates (*i*.*e*. with probability proportional to Δ*t* in a short interval [*t, t* + Δ*t*]). In our simulations, however, we use a *τ* -leap approximation where we perform all reactions for an interval of length *τ* before updating the propensity functions.

| Event | Jump | Rate |
| --- | --- | --- |
| | $(\Delta S^A, \Delta I_x^A)$ | |
| Birth | $(1, 0)$ | $b\mathcal{A}$ |
| Infection | $(-1, 1)$ | $(\beta_x I_x^A S^A)/\mathcal{A}$ |
| Death | $(-1, 0)$ | $dS^A$ |
| | $(0, -1)$ | $(d + \alpha + Uf)I_x^A$ |
| Mutation | $(0, -1)$ | $U(1 - f)I_x^A$ |
| | $(0, +1)$ | $U(1 - f)\sum_{\mathbf{u}} I_{\mathbf{x}-\mathbf{u}}^A \rho(\mathbf{u})$ |

**Figure S1:**
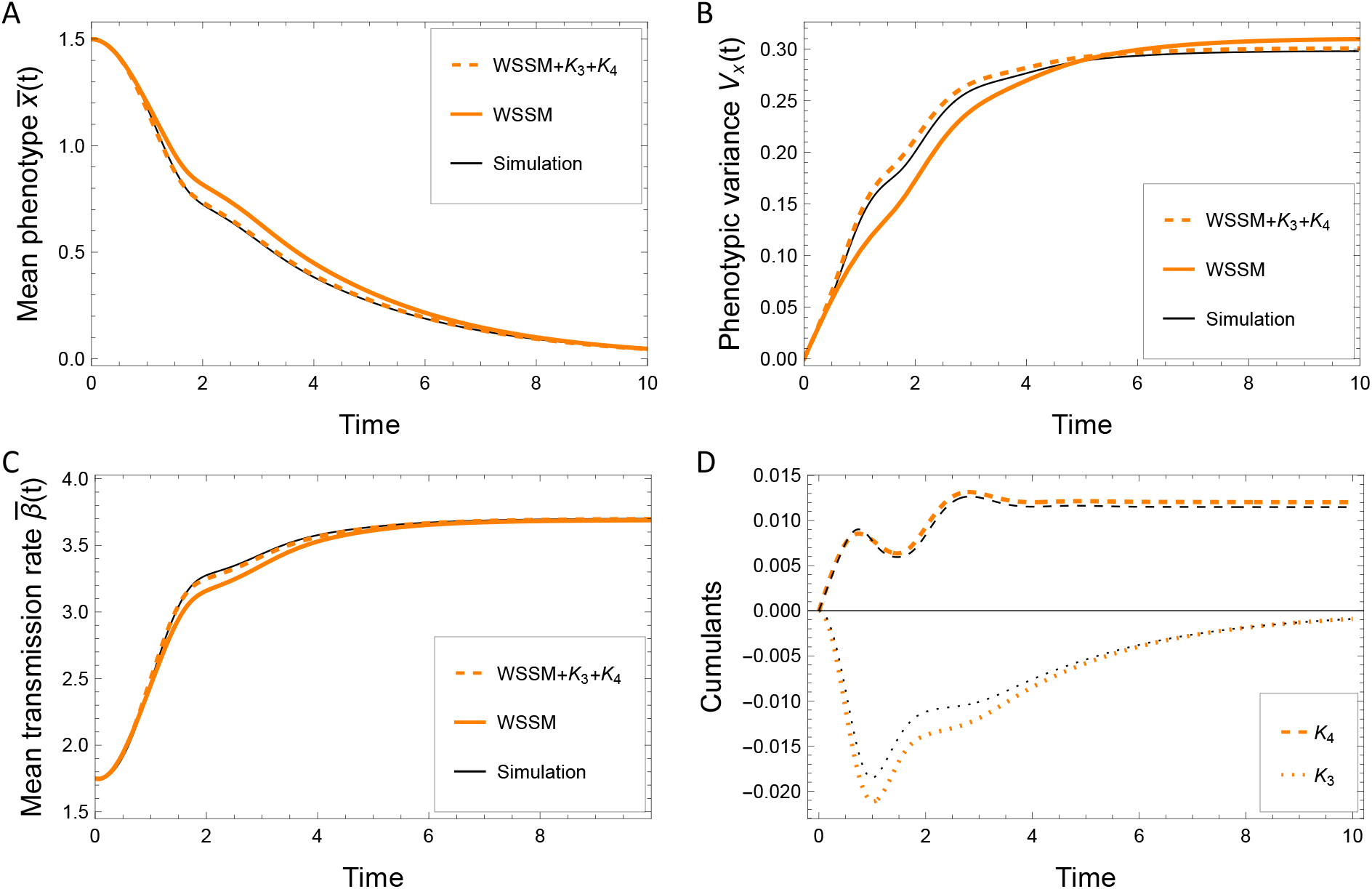
Accounting for higher moments of the mutation distribution increases the phenotypic variance transiently (which yields faster rates of viral adaptation), but it reduces the long-term phenotypic variance (which yields lower load of mutation. *L*_*M*_ ). The evolutionary dynamics are shown according to the WSSM model (orange) or taking into account mutations of larger effect through the addition of cumulants *K*_3_ and *K*_4_ (dashed orange). The result from the deterministic numerical simulations are shown in black. We show (A) the mean phenotype 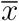,(B) the phenotypic variance *V*_*x*_ and (C) the mean transmission rate 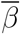.In (D) these third (dotted line) and fourth cumulants (dashed line) of the genetic distribution are shown for the WSSM+cumulants approach in orange, and for numeric simulations in black. The parameter values used are: *b* =4, *d* =1, *β*_0_= 4, *α* =1, *s*_*β*_ =1, *U*= 4, *f* =0.4, *λ* =0.1, *n*= 2. Initial conditions were 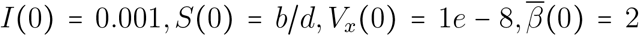.For the deterministic simulations using a grid of phenotypes, the additional parameters were *l* = 120, *x*_*max*_ = 3, *dt* = 0.02, *t*_*max*_ = 10.

**Figure S2:**
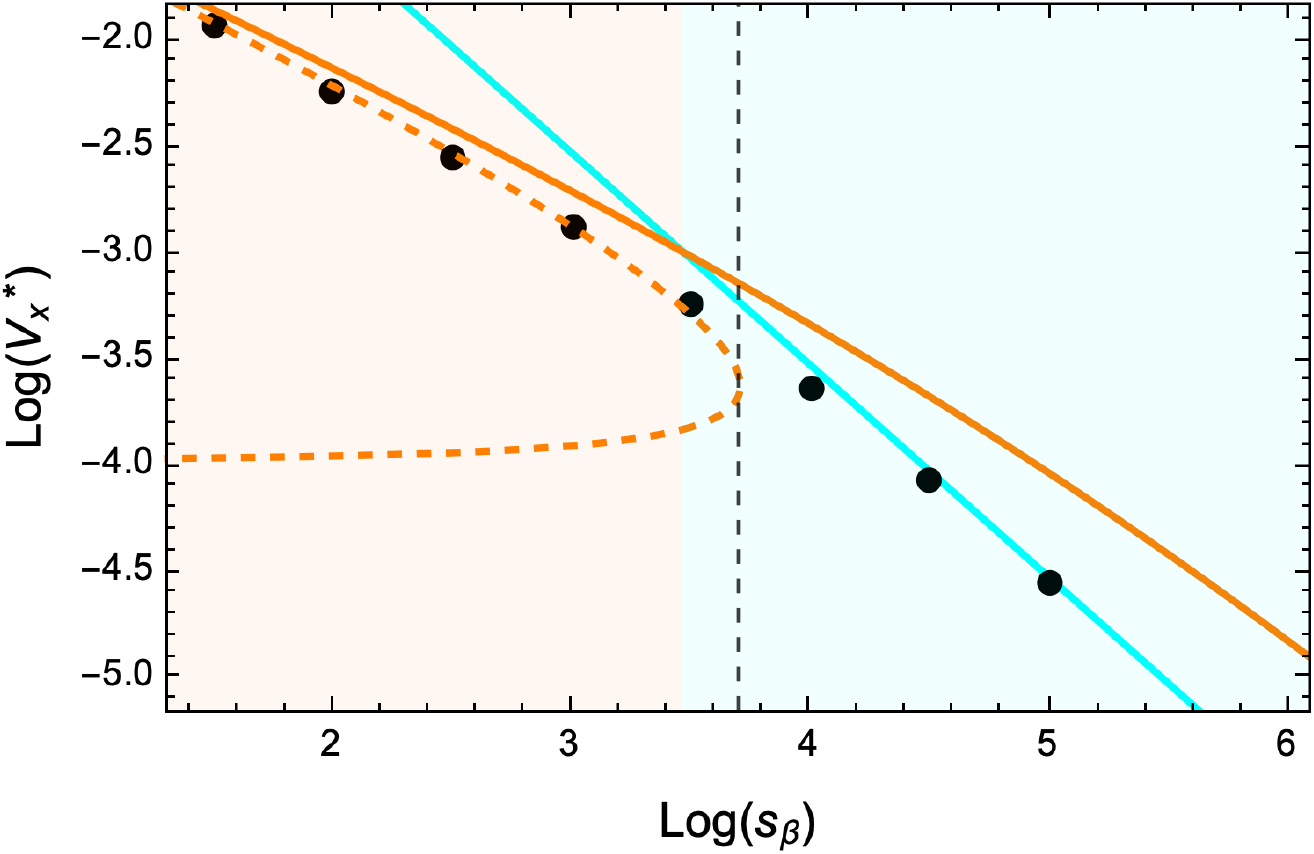
Comparison of the equilibrium variance under different regimes. The equilibrium variance *V*_*x*_ is shown as a function of the selection parameter *s*_*β*_ in a log-log scale. The results are shown for three derivations: the WSSM approximation in orange, the results from the cumulant approach up to *K*_4_ in Dashed orange and the “House of Cards” (HC) approximation in cyan. The shaded orange and cyan areas indicate the validity conditions of the WSSM and the House of Cards regimes, respectively (see equation (S.52)). The vertical dashed line shows the maximum value of *s*_*β*_ for which the WSSM+cumulant approach is valid. The parameters used were: *b* =3, *d* =1, *β*_0_= 4, *α*= 1, *U* = 4, *f*= 0.4, *λ* =0.05, *n*= 2. For the deterministic simulations using a grid of phenotypes, the additional parameters were 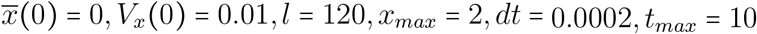.

**Figure S3:**
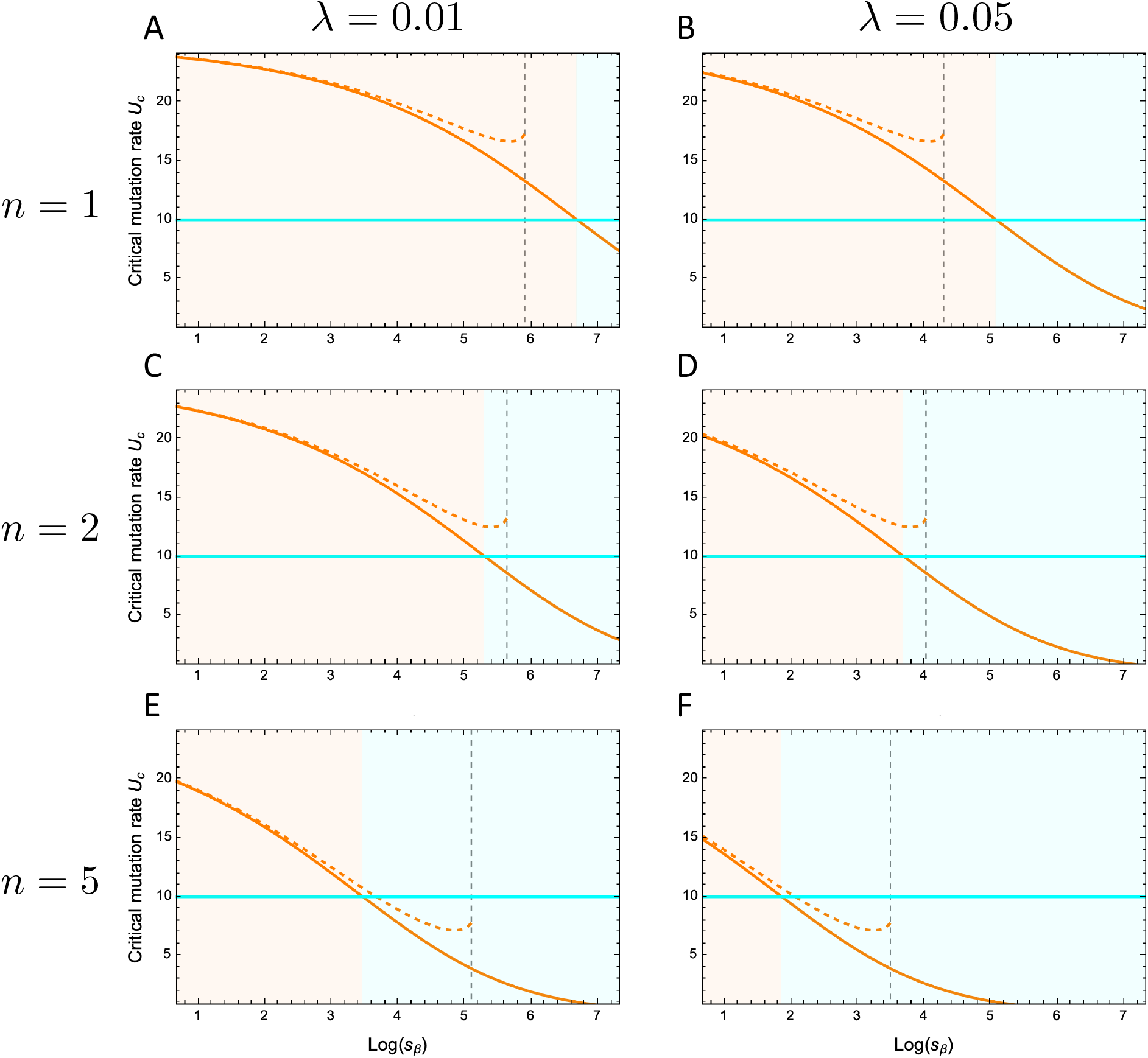
Critical mutation rate for lethal mutagenesis under different regimes. The critical mutation rate *U*_*c*_ is shown as a function of the selection parameter *s*_*β*_ for different values of mutational variance *λ* and phenotypic dimensions *n*. The results are shown under three approximation: the WSSM approximation (orange), the approximation accounting for *K*_3_ and *K*_4_ cumulants (dashed orange), the House of Cards (HC) approximation (cyan). The shaded orange and cyan areas indicate the validity conditions of the WSSM and the House of Cards regimes, respectively (see equation (S.52)). The vertical dashed line shows the maximum value of *s*_*β*_ for which the WSSM+cumulant approach is valid. Parameters used: *b* =3, *d*= 1, *β*_0_= 4, *α*= 1, *f*= 0.4. We use *λ*= 0.01 for (A,C,E), *λ* =0.05 for (B,D,F), *n*= 1 for (A,B) and *n* =2 for (C,D), *n* = 5 for (E,F).

**Figure S4:**
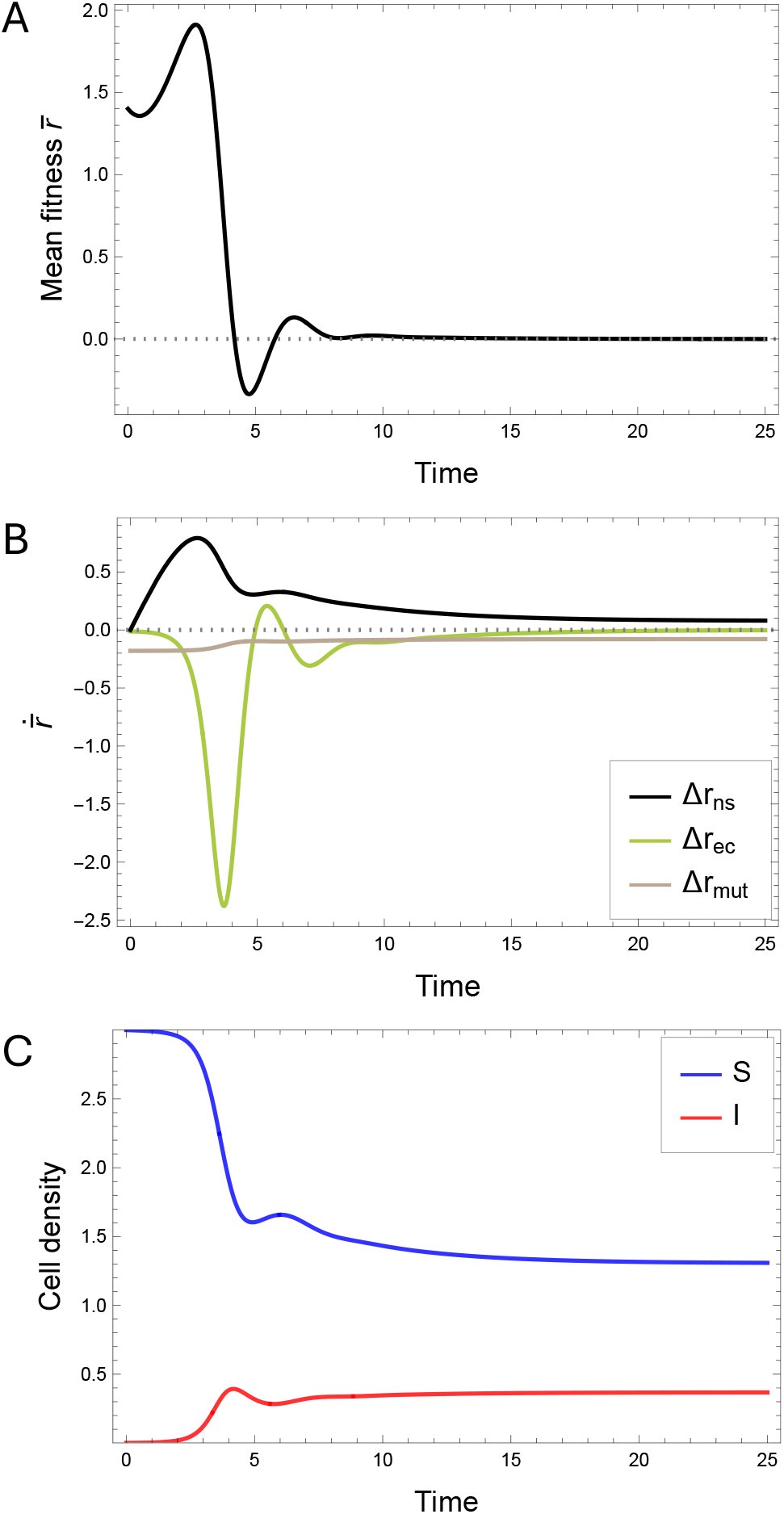
The within-host dynamics of viral adaptation. We show the epidemiological dynamics of the system starting from a clone (*V*_*x*_ (0)≪ 1). Mean fitness *r* is shown through time (a) and the different components of its dynamics are represented in (b). The black line in (b) shows the effect of natural selection (Δ*r*_*ns*_), the brown line shows the effect of mutation (Δ*r*_*mut*_) and the green line represent the feedback from the demography of the susceptible cells i.e. the environmental change (Δ*r*_*ec*_). The dynamics of the density of infected and susceptible cells are shown in (c). The parameters were *b* = 3, *d* = 1, *β*_0_ = 4, *α*= 1, *s*_*β*_= 1, *U* = 4, *f* = 0.4, *λ*= 0.005, *n*= 10. Initial conditions were 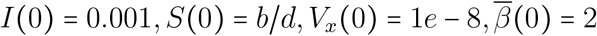

**Figure S5:**
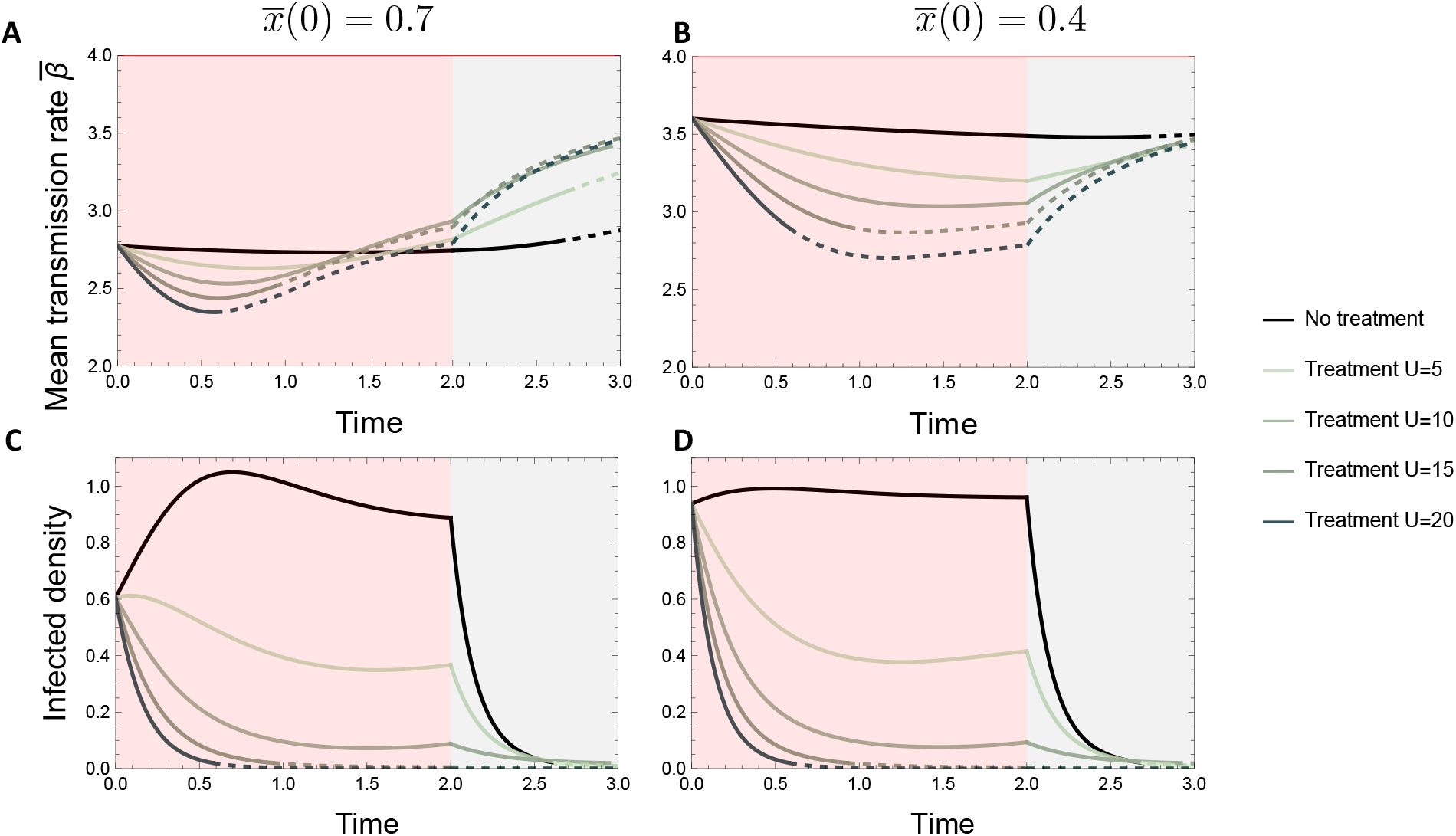
Effect of a mutagenic treatment on acute infections. The mean transmission rate (A,B) and infected density (C,D) are shown through time depending on an initial distance to the optimum of the pathogen of 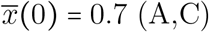 or 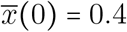 (B,D) infection, for the WSSM+cumulant model. From the start the mutation rate is increased from 1 to the value of *U* specified on the right of the plots (red shaded area). At time *t* = 2 the acute phase starts (shown in grey) and the virulence is 8-fold and so the infection is cleared even without a treatment (black curve). Simulations are initialized with clones (null variance) and the densities of susceptible and infected cells are initialized with the equilibrium values obtained with *U*= 1. The computed deterministic critical mutation rate before virulence is increased is *U*_*c*_ = 17.4. When the Infected density drops below an arbitrary value of *ϵ* = 0.02, the corresponding section of the Infected density and mean transmission rate curves become dashed. This highlights how the changes in mean transmission rate observed in these time frames are highly dependent on escaping stochastic extinction when density is low. The horizontal red line in (A,B) shows the maximum transmission rate *β*_0_. The parameters used were: *b* = 3, *d* = 1, *β*_0_ = 4, *α* = 1, *α*_*acute*_ = 8, *f* = 0.4, *λ* = 0.05, *n* = 5.

**Figure S6:**
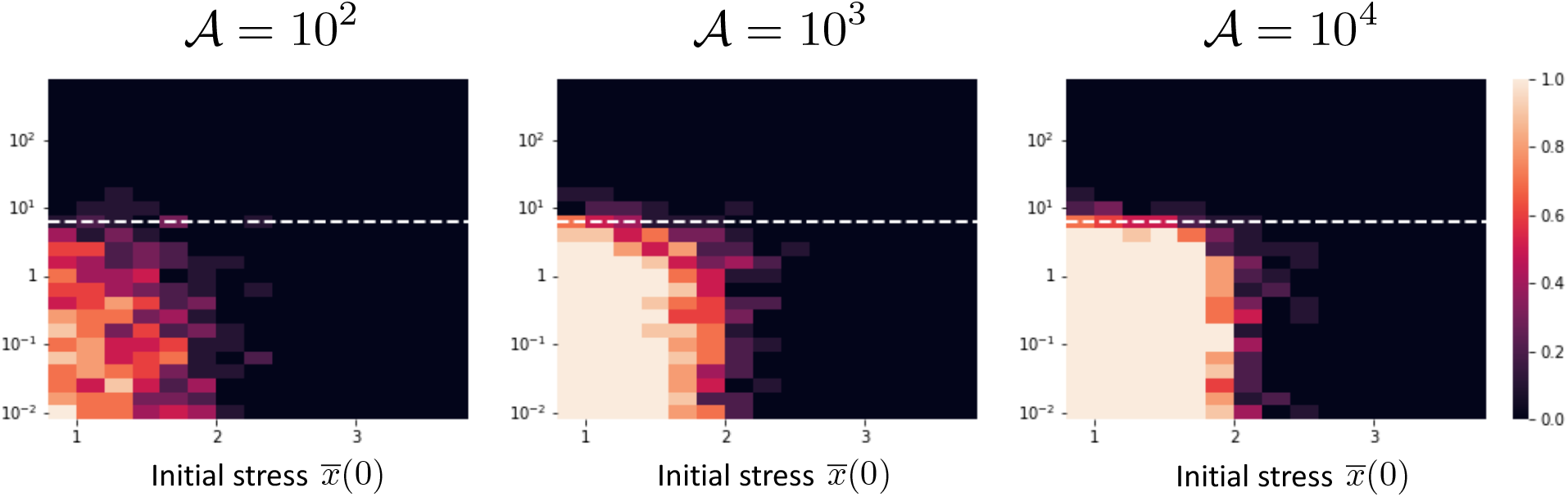
Effect of the population size on the probability of evolutionary rescue. In each panel (A-C) the population size scaling factor 𝒜 is different and indicated on top of the plot. The simulations are initialized with the equilibrium values of *S,I* and variance *σ* for a mutation rate of *U*= 10^−2^. The color scale represent the proportion of simulations in which the infected populations survived. There are 10 simulations per parameter combination. The horizontal dashed white line represents the value of critical mutation rate *U*_*c*_ above which the infected population goes to extinction in the deterministic model. The parameters used were: *b* = 2, *d*= 1, *β*_0_= 4, *α* = 4, *λ* = 0.05, *n* = 2, *s*_*β*_ = 0.5, *τ* = 0.05, *t*_*max*_ = 50.

**Figure S7:**
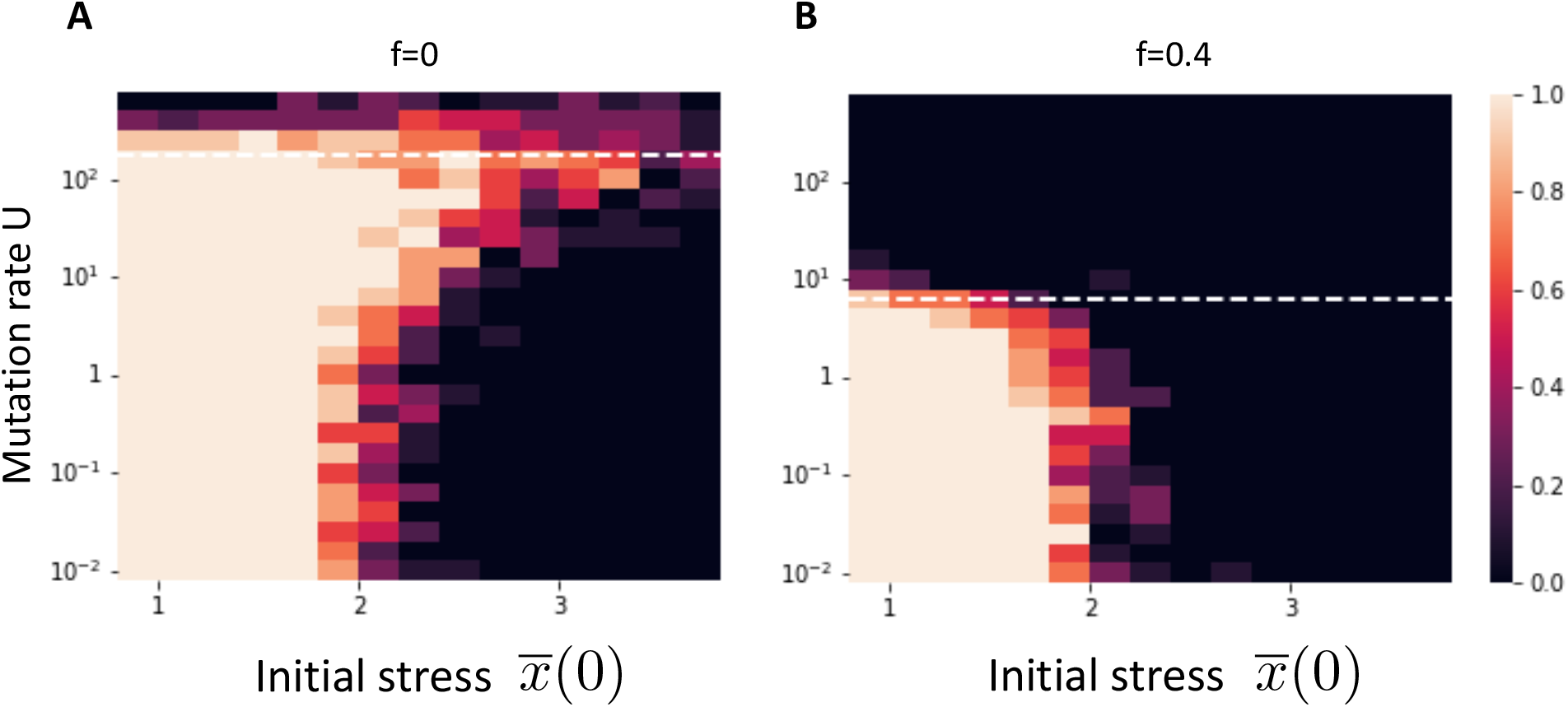
Probability of evolutionary rescue in the absence of demographic feedback, with (A) and without (B) lethal mutations. In these simulations, the density of susceptible is constant and equal to *b/ d*. The simulations are initialized with the equilibrium values of *S,I* and variance *σ* for a mutation rate of *U*= 10^−2^. The color scale represent the proportion of simulations in which the infected populations survived. There are 10 simulations per parameter combination. The horizontal dashed white line represents the value of critical mutation rate *U*_*c*_ above which the infected population goes to extinction in the deterministic model. The parameters used were: *b* = 2, *d* = 1, *β*_0_ = 4, *α* = 4, *λ* = 0.05, *n* = 2, *s*_*β*_ = 0.5, A = 1000, *τ* = 0.05, *t*_*max*_ = 50.

**Figure S8:**
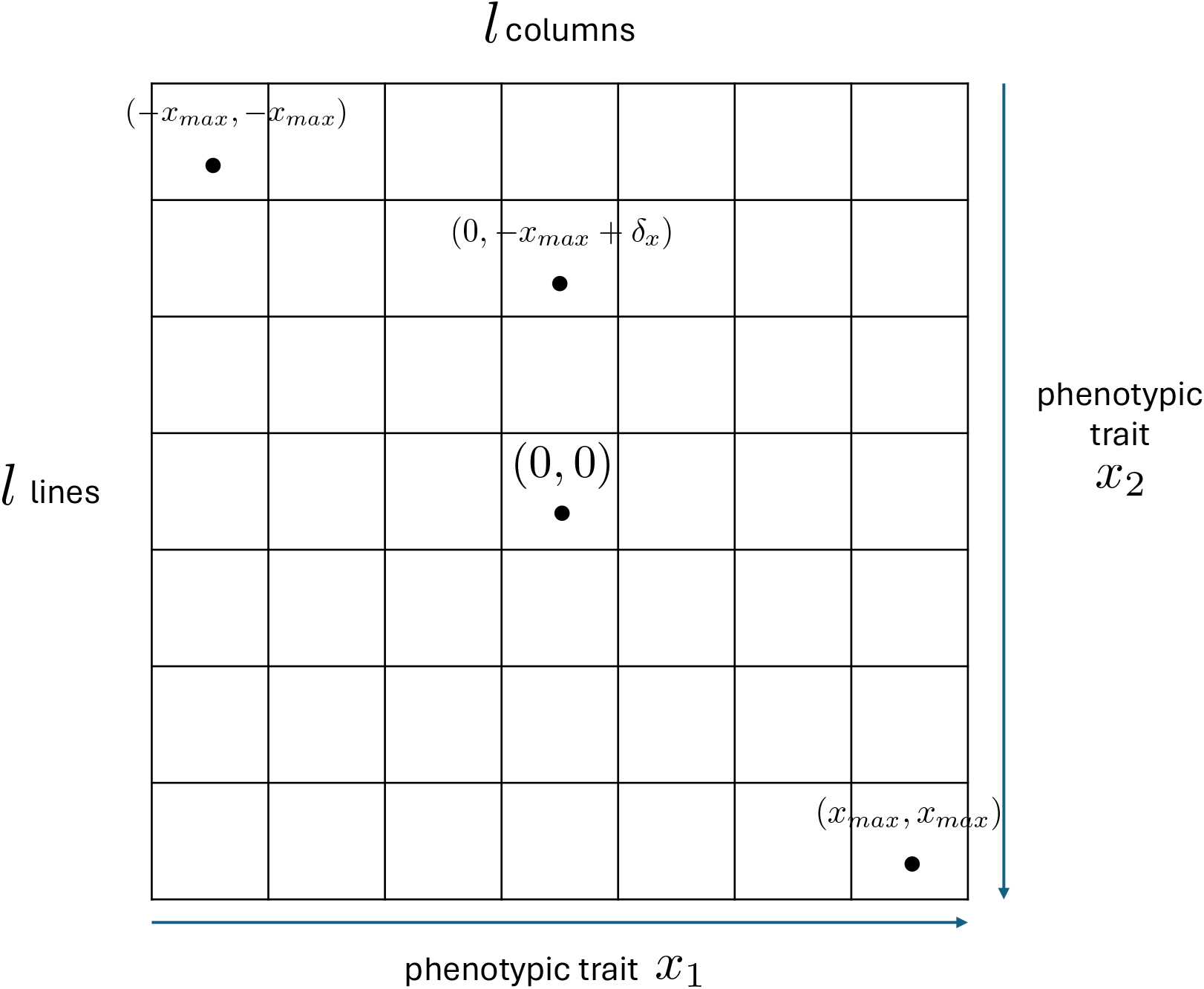
Grid of phenotypes for numeric simulations. The grid is of size *l× l* and spans phenotypes from (−*x*_*max*_, *x*_*max*_) to (*x*_*max*_, *x*_*max*_). with a phenotype step size of *δ*_*x*_. Example phenotype values are represented on the grid.

**Figure S9:**
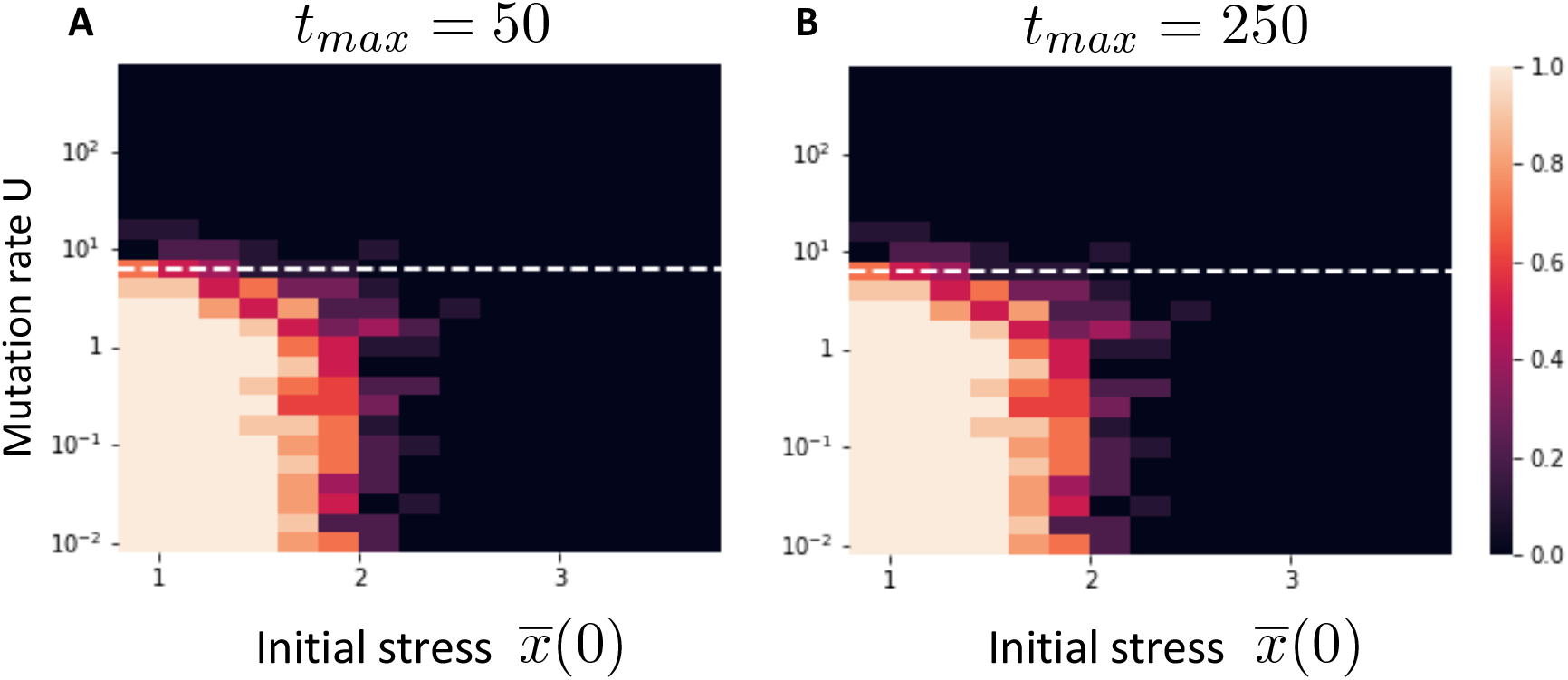
Effect of the duration of the simulation on the probability of evolutionary rescue. The simulations are initialized with the equilibrium values of *S,I* and variance *σ* for a mutation rate of *U =*10^−2^. The color scale represent the proportion of simulations in which the infected populations survived. There are 10 simulations per parameter combination. The horizontal blue line represents the value of critical mutation rate *U*_*c*_ above which the infected population goes to extinction in the deterministic model. The parameters used were: *b=* 2, *d =*1, *β*_0=_ 2.5, *α=* 1, *λ=* 0.05, *n* = 2, *η* = 1000.

## Notes

### Competing Interest Statement

The authors have declared no competing interest.

### Summary of Updates

This version of the manuscript has been revised to add the prediction of the critical mutation rate for a selection of viruses, as well as to enhance the overall presentation of the results.

